# Acoustic camera system for measuring ultrasound communication in mice

**DOI:** 10.1101/2021.12.12.468927

**Authors:** Jumpei Matsumoto, Kouta Kanno, Masahiro Kato, Hiroshi Nishimaru, Tsuyoshi Setogawa, Choijiljav Chinzorig, Tomohiro Shibata, Hisao Nishijo

## Abstract

To investigate molecular, genetic, and neural mechanisms underlying social behaviors and their deficits in neuropsychiatric disorders, social communication via ultrasonic vocalizations (USVs) in mice has received considerable attention as a powerful experimental model. The advances in sound localization technology have facilitated the analysis of vocal interactions between multiple mice. However, existing sound localization systems are built around distributed-microphone arrays, which require a special recording arena and long processing time. Here we report a novel acoustic camera system, USVCAM, which enables simpler and faster USV localization and assignment. The system comprises recently developed USV segmentation algorithms with a modification for overlapping vocalizations that results in high accuracy. Using USVCAM, we analyzed USV communications in a conventional home cage, and demonstrated novel vocal interactions in female ICR mice under a resident-intruder paradigm. The extended applicability and usability of USVCAM may facilitate future studies investigating normal and abnormal vocal communication and social behaviors, as well as the underlying physiological mechanisms.

## Introduction

Ultrasonic vocalizations (USVs) are used for communication by many rodent species (Sales, 2010). Recently, USV communication in mice has received considerable attention as a powerful experimental model to investigate molecular, genetic, and neural mechanisms underlying social behaviors and deficits (Fischer and Hammerschmidt, 2011; Lahvis et al., 2011; Konopka and Roberts, 2016). Since differences in acoustic features of USVs are insufficient for discriminating individuals (Goffinet et al., 2021) and USVs are not associated with visually distinctive movements (e.g., opening mouth), it has not been feasible to identify which mouse in a group emits a certain USV. Therefore, USV communication has been left unexplored in most studies on social behavior in mice, despite its importance. Recent advances in sound localization technology in these studies have greatly facilitated the analysis of vocal interactions between multiple subjects (Neunuebel et al., 2015; Sangiamo et al., 2020). However, to date, sound localization systems for mouse USVs (Neunuebel et al., 2015; Heckman et al., 2017; Warren et al., 2018) are built around distributed-microphone arrays (Fig. 1a; distributed-microphone systems), which require a special recording arena that is often equipped with reticulated (acoustic transparent) walls, surrounded by multiple microphones to investigate aspects of animal behavior. The distributed-microphone systems are also computationally expensive, practically requiring the use of a computer cluster for data processing (Warren et al., 2018). These technical requirements present a major obstacle for the application of such sound localization technology in established behavioral paradigms in many laboratories. To facilitate the application of sound localization technology in commonly used behavioral paradigms, we developed a novel system, named USVCAM (Fig. 1b), which was inspired by the acoustic camera, a portable device that combines a camera and a compact microphone array to visualize sound sources on camera images. In addition to the recording simplicity, the processing speed in this system is considerably faster than that of the distributed-microphone system, since the computation time window is much smaller (Fig. 1b). USVCAM also achieves more accurate localization by using a recently developed high accuracy USV segmentation algorithm (Tachibana et al., 2020) to reduce noise and focus on the USV segments that require processing. Furthermore, the segmentation algorithm was modified to discriminate overlapping vocalizations from multiple subjects. Finally, we demonstrate the performance and effectiveness of USVCAM by analyzing USV communication in a home cage, which is challenging to perform with distributed-microphone systems.

**Figure 1.**
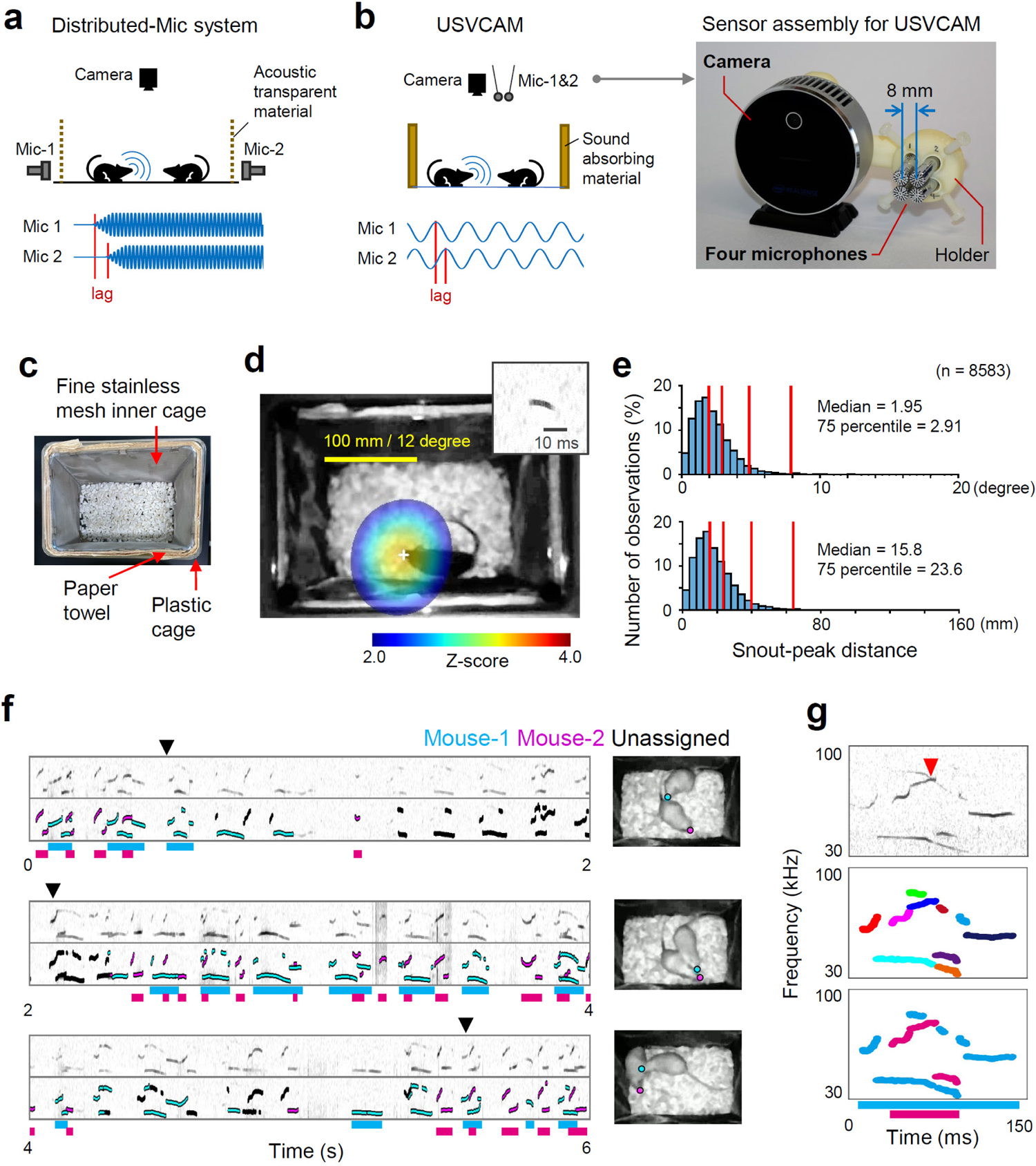
USV localization and assignment using USVCAM. **(a)** A schematic showing the setup (top) and the signals received from the microphones (bottom) using the distributed-microphone (Mic) system. Only two microphones are shown for simplicity. Because the resolution of sound localization depends on the time lags of the sound arrival, microphones are located on the sides of the recording chamber, with acoustic transparent walls to maximize the lags. **(b)** A similar schematic (left) and a picture of the sensor assembly (right) of the USVCAM. Because USVCAM utilizes phase lags of sound waves for sound localization, the microphone array can be set in one place. **(c)** A home cage equipped with the custom inner cage. **(d)** An example of sound localization of a USV segment (inset). The white cross signifies the peak of the spatial spectrum. **(e)** Distributions of the localization errors are shown in degrees (top) and millimeters (bottom). Red vertical lines indicate 50th, 75th, 95th, and 99th percentiles of the distributions, respectively. The error distributions separately calculated for B6 and ICR mice are shown in Supplementary Fig. 2. **(f)** An example of a USV assignment. The original spectrograms and those overlayed with the assignment results are shown. Bars under the spectrograms indicate syllables assigned to each mouse. The video frames at the black arrows are shown on the right. The snout positions are labeled by colored circles. **(g)** An example of the segmentation and assignment of the overlapping USVs emitted from different mice. Top, spectrogram; middle, the segmentation result (different colors indicate different segments); bottom, the assignment result. The frequency (y-axis) ranges of all spectrograms in the figure are 30 to 100 kHz.

## Results and Discussion

Two strains of mice (C57BL/6 [B6] and ICR), which have distinct vocal communications and social behavior characteristics (Asaba et al., 2014; Golden et al., 2017) were used in the following experiments. First, to validate the performance of USV localization and assignment (i.e., to identify which animal vocalizes), we recorded the vocalizations from a single male mouse exploring an empty home cage of a female mouse (single mouse experiment), in which we were certain about the source of each USV. To achieve accurate sound localization, we reduced sound reflection by inserting disposable paper towels between a conventional plastic cage and a custom inner cage made of fine stainless mesh (Fig. 1c and Supplementary Fig. 1). Fig. 1d shows an example of the spatial spectrum of a detected USV segment and the estimated source location (the peak of the spectrum) in a single mouse experiment (Supplementary Videos 1 and 2; see Methods for details of the algorithm). The median errors of localizations were 1.95 degrees (direction from the microphone array) and 15.8 mm (Fig. 1e), which are comparable to those reported for previous systems (Neunuebel et al., 2015; Heckman et al., 2017; Warren et al., 2018). Furthermore, the localization process for each USV segment took approximately 0.1 s using a single consumer personal computer (PC), which is more than 1000 times faster than the computation time (6 min/segment) reported in a previous distributed-microphone system (Warren et al., 2018).

In USVCAM, each USV segment is assigned to an individual whose average power at the snout position is significantly higher than that of other mice. We tested the assignment precision by performing simulations using data from the single mouse experiment with one to three virtual mice added at random locations within the cage (Supplementary Table 1). The precisions were around 99%, which is consistent with the confidence threshold (0.99) used for assignment. We also counted the assigned segments during social interaction experiments between two or three real mice (Supplementary Table 2). Despite the close interaction in the home cage, around half or more segments were assigned in most cases, which is comparable with the performance of distributed-microphone systems (Neunuebel et al., 2015; Warren et al., 2018). The assignment ratios in ICR mice were relatively smaller than those in B6 mice because snout locations were sometimes unavailable when the snouts were hidden under another mouse during interactions between ICR mice; moreover, the ICR mice interacted closely more frequently. Fig. 1f shows an example of an assignment during the interaction between a pair of female ICR mice. Most USV segments were assigned to one of the mice, except for when the snouts of the mice were very close (see also Supplementary Videos 3 and 4). To analyze the vocal patterns, the assigned segments were finally integrated into syllables based on the gaps between segments (bars under the spectrogram in Fig. 1f and g). USVCAM separates segments around crossing points (red arrow head in Fig. 1g; Supplementary Fig. 2), which helped discriminate overlapping USVs from different subjects (Supplementary Fig. 3). The ability to segment and localize overlapping USVs is another novel aspect of this system.

Finally, we evaluated the effectiveness of USVCAM in an actual behavioral experiment by using it to analyze USV communications under the resident-intruder (R-I) paradigm, which has been challenging with previous systems (Fig. 2). All mice were tested with both a female (vs-F) and a male (vs-M) as both resident (R) and intruder (I). Fig. 2a shows the mean call rates (number of assigned syllables/min) during each type of session (see Supplementary Table 3 for statistical results). Interestingly, ICR females exhibited USVs even when they interacted with a female as an intruder and when they interacted with a male as a resident. However, it has previously been reported that in other strains the primary sender of vocalizations is the resident and the male during interactions between the same and different sex, respectively, in experiments using devocalization or anesthetization (White et al., 1998; Holy and Guo, 2005; Hammerschmidt et al., 2012). Furthermore, we compared the call rates during different actions of the self and other in the female ICR group, in which vocal interactions were most frequently observed (Fig. 2b and Supplementary Table 4). This result indicates that although the call rate changed depending on the ongoing action, as reported previously (Sangiamo et al., 2020), the call rate was also modulated by the social context, even during the same types of actions. Finally, we analyzed acoustic features of USVs using dimension reduction by a variational autoencoder (unsupervised learning methods; Goffinet et al., 2021) to determine whether the distribution of acoustic features (i.e., the vocal repertoire) in ICR mice changes depending on social contexts (Fig. 2c and Supplementary Figs. 4 and 5).

**Figure 2.**
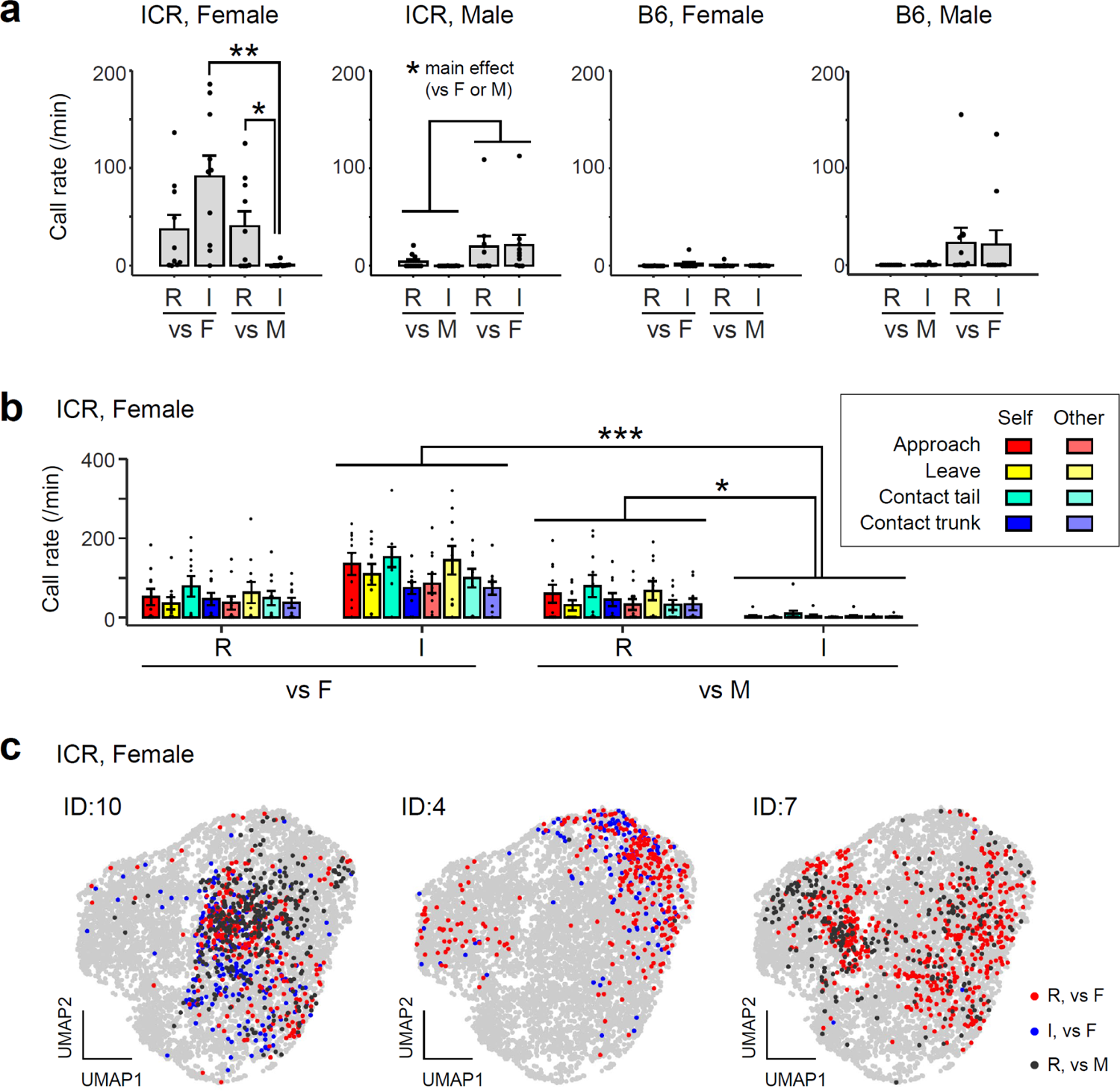
Analysis of mouse pair interactions under the resident-intruder (R-I) paradigm using USVCAM. (**a**) Comparisons of call rates (number of assigned syllables per minute) of subjects in different social contexts. R, the subject was a resident; I, the subject was an intruder; vs F, the partner was a female; vs M, the partner was a male. Each dot represents an individual mouse. Error bars, standard error of the mean (s.e.m.); \*\**p* < 0.01, \**p* < 0.05, simple main effects analysis. (**b**) Call rates of female ICR mice during different actions by the subjects (self) and partners (other). See Supplementary Figure 15 for the definition of the actions. Each dot represents an individual mouse. Error bars, s.e.m.; \*\*\**p* < 0.001, \**p* < 0.05, simple main effects analysis. (**c**) UMAP projection of the acoustic features of syllables extracted using the VAE (Goffinet et al., 2021). Examples of three ICR female mice are shown (ID, identity number of the mouse). Each point represents an assigned syllable. Red, blue, and black indicate the syllables of the subject recorded in different sessions. Gray points are all the other syllables recorded during the pair interaction experiments.

Results revealed no significant changes in the vocal repertoire depending on social contexts, although the vocal repertoire was dependent on individuals, as previously reported (e.g., Goffinet et al., 2021). To the best of our knowledge, this is the first report conducting a systemic analysis of complex acoustic features of syllables, including those that overlap. Taken together, the results of the USVCAM application revealed a novel characteristic of vocal communication in ICR mice under the resident-intruder paradigm. In summary, we developed an acoustic camera system, USVCAM, which enables simple and fast USV localization and assignment. The system incorporates a recently developed USV segmentation algorithm with a modification that permits discrimination between overlapping vocalizations to achieve high accuracy. We applied USVCAM to analyze USV communications in home cages. Home cage recording is a popular method used in many experimental paradigms because it allows the observation of undisturbed behavioral expression and thus is ideal for investigating important aspects of social behavior (e.g., resident-intruder, sexual, and mother-infant interactions; Kikusui, 2013). However, previous systems have found this challenging using a conventional home cage. USVCAM revealed novel characteristics of vocal communication between ICR mice, suggesting that it is effective in characterizing the social behaviors of various mice strains, such as those that have been genetically modified to establish disease models. USV is an important social signal in rodents (Sales, 2010), and previous studies have reported that USVs from different subjects (especially self and others) have different effects on brain activity and behavior (Rao et al., 2014; Matsumoto et al., 2016; Neunuebel et al., 2015; Sangiamo et al., 2020). Thus, sound source localization is fundamental for studying the dynamics of social behavior and its underlying mechanisms, as well as behavior phenotyping. Taken together, the extended applicability and usability of USVCAM may facilitate future studies investigating normal and abnormal vocal communication, social behaviors, and the underlying molecular, genetic, and neural mechanisms.

## Methods

### Recording

The sensor assembly (Fig. 1b right) consisted of four ultrasound microphones (TYPE 4158N, ACO, Tokyo, Japan), a video camera (RealSense L515, Intel, CA, USA), and a custom three-dimensional (3D)-printed holder for the sensors. A square phased microphone array (around 8 mm on each side) was composed with the holder. The distance between the camera and the center of the microphone array was approximately 54 × 9 × 6.5 (depth) mm (microphones were placed in front of the camera). The audio data was captured by each microphone, amplified with a four-channel microphone amplifier (BSA-CCPMA4-UT20, Katou Acoustics Consultant Office, Kanagawa, Japan), and sampled at 384 kHz using an analog-digital converter (PCIe-6374, National Instruments, TX, USA). An infrared image (resolution: 640 × 480 pixels; field of view: 70° × 55°) was captured with the camera at 30 Hz. The audio and video data were stored on the same PC (Elitedesk 800 G5 TW, Hewlett-Packard Inc., CA, USA) with timestamps of each data for audio-video synchronization, using a custom recording software written in Python (Python Software Foundation, NC, USA).

Reducing sound reflection in recording environments is crucial for ensuring accurate sound source localization based on the time/phase lags (Fig. 1b). To this end, recordings were performed in a soundproof box (63 × 53 × 84 [height] cm) with 20 mm thickness sound-absorbing melamine foam on the walls and ceiling (Supplementary Fig. 6). We also designed a custom inner cage for home cage recording (Fig. 1d and Supplementary Fig. 7). The inner cage was made from fine stainless mesh (mesh size: 150 mesh/inch). Clean disposable paper towels (Prowipe, Daio Paper Corp, Tokyo, Japan) were inserted into the space (approximately 5 mm) between the inner cage and the plastic home cage to suppress the influence of sound reflection (see Supplementary Fig. 1 for the effect of the inner cage).

### USV segmentation

For the segmentation (detection) of the USVs in the audio data, the USVSEG algorithm (Tachibana et al., 2020) was used with slight modifications. The USVSEG algorithm can robustly segment USVs from background noise by generating a stable spectrogram using the multitaper method and flattening the spectrogram by liftering in the cepstral domain. Although several different algorithms have been proposed for USV segmentation, a recent benchmarking study indicated that the USVSEG is comparable to the state-of-art method (Fonseca et al., 2021). We modified the USVSEG algorithm to separate crossing USVs emitted from different animals (Fig. 1g and Supplementary Fig. 8). First, a binary image of the spectrogram peaks was created (pixel size = 0.5 ms × 750 Hz) using the peaks obtained using the original USVSEG algorithm (Supplementary Fig. 8c). Second, the binary image was dilated twice and eroded once with a 3 × 3 square structuring element to connect the spatially neighboring components using imdilate() and imerode() functions in MATLAB (Mathworks, MA, USA; Supplementary Fig. 8d). Third, the corner points (i.e., the crossing points and edges) were detected using the corner() function in MATLAB (Supplementary Fig. 8e), and the pixels within the rectangles (3 × 15 pixels) centered on the corners were erased to cut segments at the crossing points. Finally, the boundaries of the subsegments were calculated by applying the watershed transform to the image using the watershed() function in MATLAB (Supplementary Fig. 8f), and the spectral peaks were grouped according to the boundaries (Supplementary Fig. 8g). Small segments with ≤ 3.0 ms were excluded from the subsequent analysis. We validated the algorithm with synthetic data, in which pairs of syllables recorded from a single mouse were overlapped (Supplementary Fig. 3). For test data generation, a pair of syllables recorded in a single mouse experiment were randomly selected, and the peaks of each syllable were detected. The peaks of the two sources were overlayed with a random time shift (± 20 ms). When the peaks from the different sources were close, one stronger peak was selected according to the USVSEG algorithm, and only one peak was selected within a narrow bandwidth. In total, 10,000 (5,000 from B6 and 5,000 from ICR mice data) overlapping syllables were generated, and the segmentation algorithms were applied. To test the effect of the modification of the segmentation algorithms, we compared the results between using the proposed algorithm and the algorithm without corner detection. To quantify the quality of the segmentation, the contamination ratio was defined as (1/*N*)×*∑n_i_*, where *n_i_* and *N* represent the number of contaminated points (i.e., the points from the minor source) in the *i*-th segment and the total number of points, respectively.

### USV localization

Using the conventional (Bartlett, or delay-and-sum) beamformer, the power (*P*) of sound arriving from a given spatial location (***r***) was calculated as follows (Krim and Viberg, 1996):

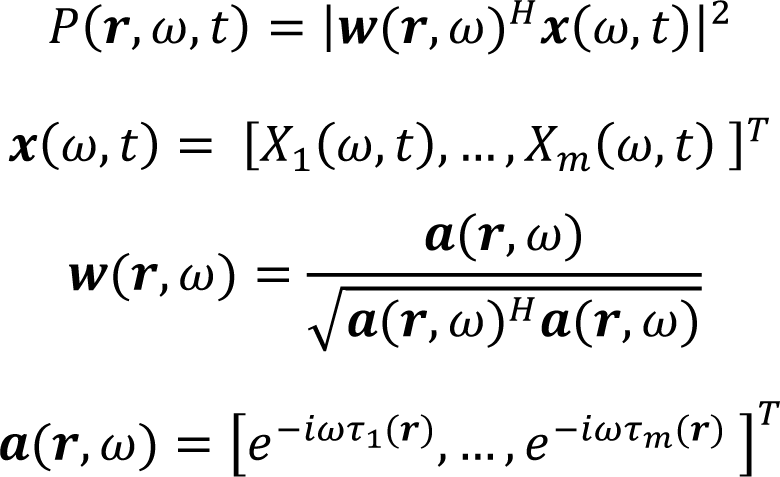

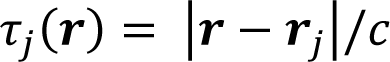

where *ω* and *t* are frequency and time indices, respectively. *X_j_(ω,t)* represents the short-term Fourier transform of the signal captured with the *j*-th (*j = 1,…, m*) microphone. ***r****_j_* is the position of the *j*-th microphone, *c* is the speed of sound, and *τ_j_* is the expected time delay of the signal arriving at the *j*-th microphone from the sound source location ***r***. *T* and *H* denote the transposition and conjugate transposition, respectively. The beamformer shifts the signals captured by the microphones to compensate for the arrival time delays (i.e., the phase lags of the signals) using a steering vector ***a***, and sums the shifted signals to calculate the power (*P*). Thus, the function *P(**r**)* is expected to be the maximum when the location ***r*** overlaps with the actual sound source and is called the spatial spectrum of the sound. Because we are not interested in the absolute power of the signal but rather in the peak locations in the spatial spectrum for sound localization, the following normalized spatial spectrum *P_norm_* was used:

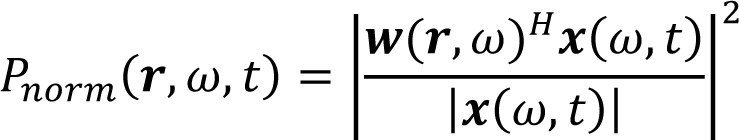

The average spatial spectrum of a given segment *P_seg_* (Fig. 1d) was subsequently defined as follows:

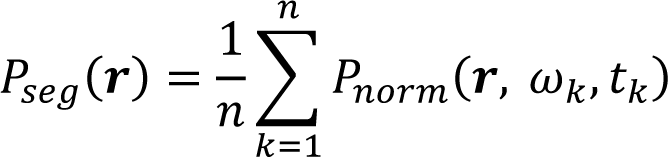

where *ω_k_* and *t_k_* represent frequency and time indices, respectively, of the *k*-th (*k=1, …, n*) peak of a USV segment in the spectrogram. To localize the USV segment, *P_seg_* was calculated at each x-y location of the camera image. The depth (z) of the sound source locations were assumed to be constant (i.e., equal to the floor of the cage). To reduce computational load, the image was binned into 5 × 5-pixel bins, and *P_seg_* were calculated for each bin. *P_seg_* outside the image were also calculated for margins of 100 pixels on each side to correctly estimate the peaks of the spatial spectrum around the edge of the camera image. The resultant spatial spectrum often showed multiple peaks in the form of a square grid (Supplementary Fig. 9) owing to the periodicity of the sound wave and the square microphone array arrangement; this phenomenon is referred to as spatial aliasing. To avoid spatial aliasing affecting the result, we positioned the microphones as close as possible (Fig. 1b) to maximize the distance between the multiple peaks in the spatial spectrum. Furthermore, we used a USV assignment algorithm that can deal with spatial aliasing (see below). The spatial spectrum was normalized with the z-score normalization to evaluate the saliency of the peaks. Peak locations (local maximums) in the spatial spectrum were searched as bins with equal values across the original spectrum and that filtered using the maximum filter (window size = 5 × 5 bins). A peak with a small height (z < 1.6, approximately < 95% in the cumulative distribution function of the standard normal distribution) was excluded from the sound source candidates. A USV segment without any clear peak (z < 2.3, approximately < 99% in the cumulative distribution function of the standard normal distribution) in the spatial spectrum was categorized as ‘unlocalized’ and was excluded from subsequent analysis. The snout-peak distance was defined as the distance from the snout of the mouse and the nearest peak in the spatial spectrum.

#### USV assignment

Supplementary Fig.10 shows an overview of the algorithm for USV assignment for each USV segment. Initially, the snout-peak distance was calculated for each mouse, and the mice with snout-peak distances that were within the distance threshold (red dotted line) were selected as candidates of the sound source (Supplementary Fig. 10a). The distance threshold was set as 99th percentiles of the distribution of the snout-peak distances in the single mouse experiments (Fig. 1e and Supplementary Fig. 10a right). If the number of candidates was zero or one, the USV segment was categorized as ‘unassigned’ or assigned to the candidate mouse, respectively. If there was more than one candidate mouse, the following test was conducted among the candidates.

To assign the USV segment to one of the candidate mice following the screening procedure above, the average power at the snout of the candidates was first calculated (Supplementary Fig. 10b). The two highest powers among the candidates were compared using a two-tailed Wilcoxon signed-rank test. The resulting *p*-value (*p*) was used for the assignment. In addition, we used the distance between the snouts of the best two mice (*d*) for the assignment because we found using a simulation that, with the same *p*-value threshold, when the snouts of the two mice become closer, more assignment errors occurred (Supplementary Fig. 11). This may have been caused by an error in sound localization itself (Fig. 1e) or an error in the video-based estimation of snout locations. Using the above two parameters *p* and *d* with the number of candidate mice (*N_c_*), the confidence for assigning the segment to the best mouse was calculated as the averaged precision of the assignment for similar conditions in the following simulation using the single mouse experiment data. In the simulation, *N_c_*-1 ‘virtual’ mice were assumed to be at random locations away from the real mouse at a distance *d*, and the precision (hit count/[hit + error counts]) of the assignment at the *p*-value threshold *p* was estimated. The simulations were performed in advance for all possible combinations of *p*, *d*, and *N_c_* (Supplementary Fig. 11), and the distribution of the precision was used as a lookup table for the confidence estimation to reduce the computational load. Since the lookup tables that had been separately calculated for B6 and ICR mice (Supplementary Fig. 12) were similar, we used a combined distribution (Supplementary Fig. 11) for assignment in this study. USV segments with a confidence level of > 0.99 were assigned to the best mice, and the others were categorized as ‘unassigned.’

The confidence estimation may be inaccurate when the two best mice are located near two different peaks in the spatial spectrum because of spatial aliasing (Supplementary Figs. 9 and 13a). In such cases, although the absolute snout-snout distance (*d*) is large, both mice should be considered good candidates. To ameliorate the problem, we calculated the distance between snouts (*d’*) after converting the snout positions into their relative positions from the nearest peaks (i.e., ‘wrapping’ the positions in a period of the spatial spectrum; Supplementary Fig. 13b) and used *d’* for the confidence estimation.

#### Merging segments into syllables

Rodent USVs consist of syllables, which have tens to hundreds of milliseconds durations with gap intervals (usually of > 30 ms). Previous studies categorizing syllable patterns reported that the patterns differed depending on the behavior, individual, and strain (Holy and Guo, 2005; Matsumoto and Okanoya, 2018; Goffinet et al., 2021). Thus, we merged the short segments (Fig. 1g) into syllables after the assignment. Specifically, segments assigned to one mouse and ‘unassigned’ segments (if they existed) with gap intervals smaller than a given threshold (a minimum gap of 30 ms) were merged into a single syllable. The assignment rate was defined as the ratio of the time-frequency points assigned to the mouse to all the points in the syllable. We only used syllables with an assignment rate of 1.0 for the behavior analysis.

#### Audible broadband vocalization detection

Mice occasionally emit audible broadband vocalizations (BBVs; i.e., ‘squeaks’) during conspecific interactions (Finton et al., 2017). BBVs are loud broadband sounds characterized by a harmonic structure. To prevent misidentifying BBVs as USVs and misassigning USVs that significantly overlap with BBVs, we excluded the ultrasound segments that overlapped with the time intervals of BBVs from the analysis. BBVs were detected automatically using the following simple algorithm. First, the recorded sound was downsampled to 38.4 kHz. Second, the spectrogram (time window = 10 ms; frequency range = 2 to 16 kHz) of the sound was calculated. Third, continuous background noise and transient (impulse-like) broadband noise were reduced by subtracting the median value of each frequency bin and the median value of each time bin, respectively. Fourth, the spectrogram was filtered using a median filter (window size = 0.5 kHz) along the frequency axis. Finally, the maximum power in the spectrogram across the frequency was calculated for each time point, and the time intervals containing BBVs were estimated as the intervals in which the maximum power exceeded a certain threshold value (we used 28 dB in this study). To check the precision of the simple BBV detector, we selected three recording sessions that involved a relatively large number of BBVs (interactions between one pair of male ICR mice and two male-female ICR mouse pairs; see below for details of the recording experiment), and a blinded experimenter compared the results of the automatic detection with the manual annotations (Supplementary Fig. 14). A total of 161 BBVs were detected in the manual annotations. Of these, 160 overlapped with the automatic detection, and one was missed by the detector. The automatic detector had only 22 additional (false-positive) detections. Thus, the results confirmed that the simple detector can effectively exclude time intervals containing BBVs from the analysis. Supplementary Table 5 shows the number of ultrasound segments that overlapped with the time intervals of BBVs in each of the social interaction experiments.

### Calibrating microphone positions

For the above USV localization algorithm, microphone positions (***r****_j_*) needed to be accurately calibrated. For the calibration, we searched the microphone positions that maximized the average power (*P_seg_*) at the snout locations for the 20 selected USV syllables in the single mouse experiment, as follows:

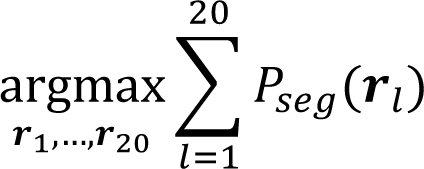

where ***r****_l_* is the snout location at which the *l*-th selected syllable was emitted. The optimization was performed using the L-BFGS-B algorithm implemented in Scipy (Virtanen et al., 2020). The syllable that emitted different parts in the recording area was selected for appropriate calibration.

#### Video tracking

USV assignment requires frame-by-frame snout locations of each mouse. USVCAM users can choose any available high-precision video tracking software (such as DeepLabCut [Lauer et al., 2021], Social LEAP Estimates Animal Poses [SLEAP; Pereira et al., 2021], and Mouse Action Recognition System [MARS; Segalin et al., 2021]) to estimate snout locations. In this study, we used AlphaTracker (Chen et al., 2020) for tracking the locations of the snout and the other body parts. The software can relatively robustly track the locations of body parts of interacting mice using deep neural networks. We prepared labeled data to train the networks by manually annotating body parts (snout, tail-base, and left and right ears) and the bounding box of the mice in 1694 and 996 randomly selected frames from the videos of B6 and ICR mice, respectively. Different networks were trained for tracking B6 and ICR mice. The outputs of AlphaTracker were manually curated to correct occasional errors (e.g., switching mouse identities and flipping snout and tail-base locations) using custom software written in Python. The resultant trajectories of the snouts were filtered using a median filter (window size = 0.16 s) and used for USV assignment. The trajectories of the bounding box, tail-base, left and right ears, and snout were filtered using a locally estimated scatterplot smoothing (LOESS) filter (time window = 0.5 s) and used for behavioral event classifications (see below). Supplementary Video 5 shows an example of the filtered trajectories used for behavioral event classification.

#### Audible sound generation from recorded USVs

For intuitive data presentation, the audible sound was created according to the results of the USV segmentation and integrated with the videos (Supplementary Videos 1–4) using a sound synthesis method proposed by Carruthers et al. (2013). First, the maximum peak of the USV segments at each time point was extracted. Then, the sound *x(t)* was generated as:

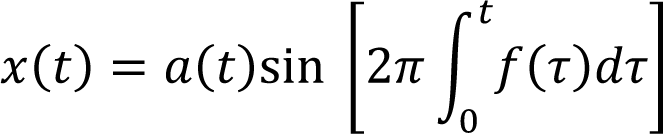

where *t* is time, and *a(t)* and *f(t)* are the amplitude and frequency of the peak, respectively. Thus, the generated sound contains no background noise. The frequency was linearly mapped from 0 to 192 kHz to 1 to 6 kHz to make the sound audible. The generated sounds were only used for visualization.

### Animals and experimental schedule

All animal experiments were performed with institutional ethics approval (from the Institutional Animal Use Committee of Kagoshima University #L21007). We used 40 adult mice: 10 males and 10 females each of the C57BL/6J (B6) and ICR strains. Mice were purchased from Japan SLC (Shizuoka, Japan) at the age of 8 weeks and housed alone in a cage (182 × 260 × 128 mm, CREA Japan, Tokyo, Japan) equipped with the custom inner cage for 1 week. Food (5L37 Rodent LabDiet EQ, PMI Nutrition International, MO, USA) and water were supplied *ad libitum*, and the animals were kept under a standard 12 h:12 h light-dark cycle. Soft paper chips were used for bedding (Japan SLC). Experiments were mainly conducted in the light phase, except for some of the recordings for system validation (the single mouse and three mice experiments, see below), which were conducted during the dark phase. The environment was maintained at a constant temperature (22–25°C) and humidity (50% ± 5%).

After a 1 week habituation period, the social interactions between mice in the home cages were recorded. On the first day, each mouse was allowed to interact with another mouse of the same sex and strain (M-M and F-F contexts), both as resident and intruder (nine mice as residents and one mouse as an intruder were tested first). On the second day, each mouse was allowed to interact with another mouse of a different sex but the same strain (M-F and F-M contexts). In these cases, all individuals were used as both a resident and an intruder, and the order in which these roles were applied to the experiment was counterbalanced. For the recording, the home cage with a resident mouse was placed in the soundproof recording box, an intruder mouse was placed in the home cage, and the behaviors of the mice were recorded for approximately 3 min. After the 2 days of paired social interaction recording, one mouse of each strain was recorded in a single mouse condition to obtain data for system validation and determine parameters for the USV assignment. In the single mouse experiment, a male mouse was placed in an empty home cage of a female mouse, and its vocalizations were recorded for approximately 10 min. We also tested the recording of a three-mouse interaction of each strain for approximately 5 min for system validation (Supplementary Table 2).

#### Data analysis

The number of syllables assigned completely (assignment rate = 1.0) was counted for each mouse for each recording. The call rate was calculated by dividing the syllable count by the recording duration. In addition, to check the relationship between USVs and specific actions, we defined the following five behavior events based on video tracking results: *approach*, *leave*, and *contact with the tail base*, *trunk*, and *snout* (Supplementary Fig. 15). Then, call rates during different behavior events were calculated separately. The call rates during contact with the snout were not analyzed because of difficulty in USV assignment when the snouts were very close. To quantify and compare the acoustic features (patterns) of syllables, we used a variational autoencoder (VAE), according to the method proposed by Goffinet et al. (2021). In this method, VAE learns to map single-syllable spectrogram images onto 32 latent features in an unsupervised manner. We reconstructed a spectrogram of an assigned syllable using the frequency and amplitude of each point in the segments of the syllable to enable the analysis of the acoustic feature of the syllables even when it temporally overlapped with the syllables emitted from the other mouse. In total, 7960 single-syllable spectrogram images were reconstructed and used for VAE training and analysis. The distribution of the syllables in latent space was visualized using Uniform Manifold Approximation and Projection (UMAP) and the difference in the distributions (vocal repertoires) between a pair of different experimental conditions was quantified using maximum mean discrepancy (MMD), according to a previous study (Goffinet et al., 2021). Distributions with fewer than 10 syllables were excluded from the MMD analysis. Statistical tests were performed using R (The R Foundation, IN, USA) and MATLAB.

## Data availability

The datasets generated and/or analyzed for the current study are available from the corresponding author on reasonable request. Part of the raw data is available in a public repository as example data to test the codes (doi:10.6084/m9.figshare.17121275).

## Code availability

The codes of the USVCAM system (for recording, localization, and assignment) are available in a public repository (https://github.com/MatsumotoJ/usvcam). Other codes used for further analysis are available from the corresponding authors on reasonable request.

## Acknowledgments

We would like to thank Ryosuke Tachibana for providing helpful explanations of the USVSEG algorithm, and Takafumi Tamura and the staff at Engineering Machine Shop, University of Toyama, for technical assistance for manufacturing the mesh inner cage and the 3D-printed holder. This work was supported by Grant-in-Aid for Scientific Research from Japan Society for the Promotion of Science (nos. 16H06534, 21K06438, 19H04912), the Firstbank of Toyama Scholarship Foundation Research Grant, Tokyo Biochemical Research Foundation, Takeda Science Foundation, Grant for Basic Science Research Projects from The Sumitomo Foundation, and RIKEN Collaborative Technical Development in Data-driven Brain Science.

## Author contributions

J.M. developed the USVCAM system. T.Sb. advised on algorithm selection. M.K. advised on the design of the microphone array. K.K. collected experimental data. J.M., K.K., C.C., T.Sg., H.Nm., and H.Nj. analyzed data. J.M., K.K., H.Nm., H.Nj., T.Sb., and T.Sg. wrote the manuscript. All authors contributed to the article and approved the submitted version.

---

**Supplementary Figure 1.**
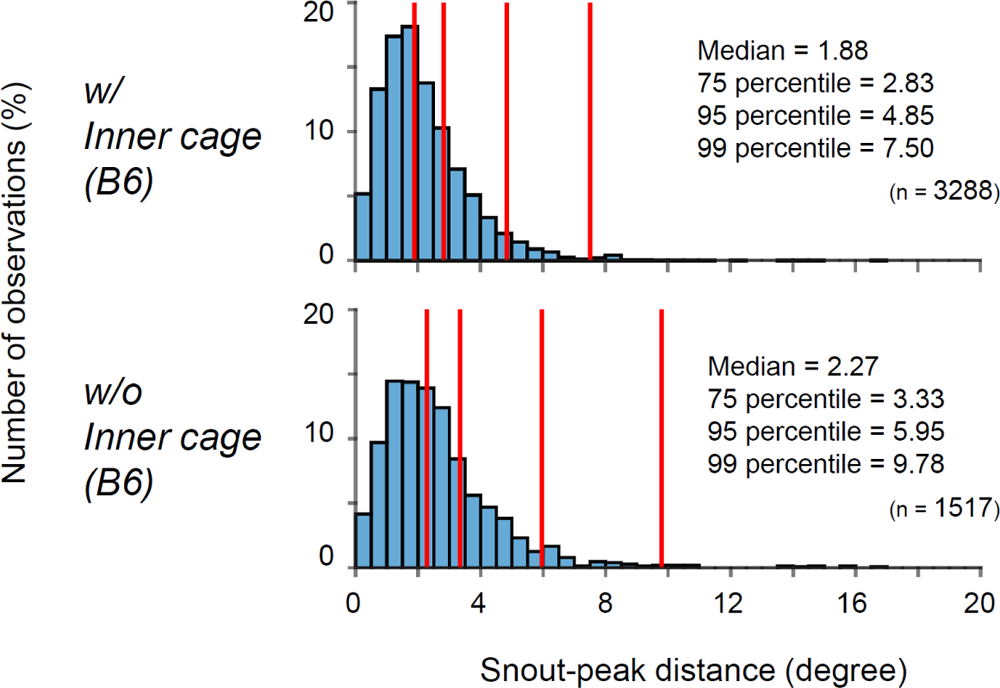
Error histograms similar to those in Figure 1e, in the home cage with (top) and without (bottom) the inner cage and paper towels in a B6 mouse. Note that more outliers were observed without the inner cage (bottom), which demonstrates the effect of the inner cage.

**Supplementary Figure 2.**
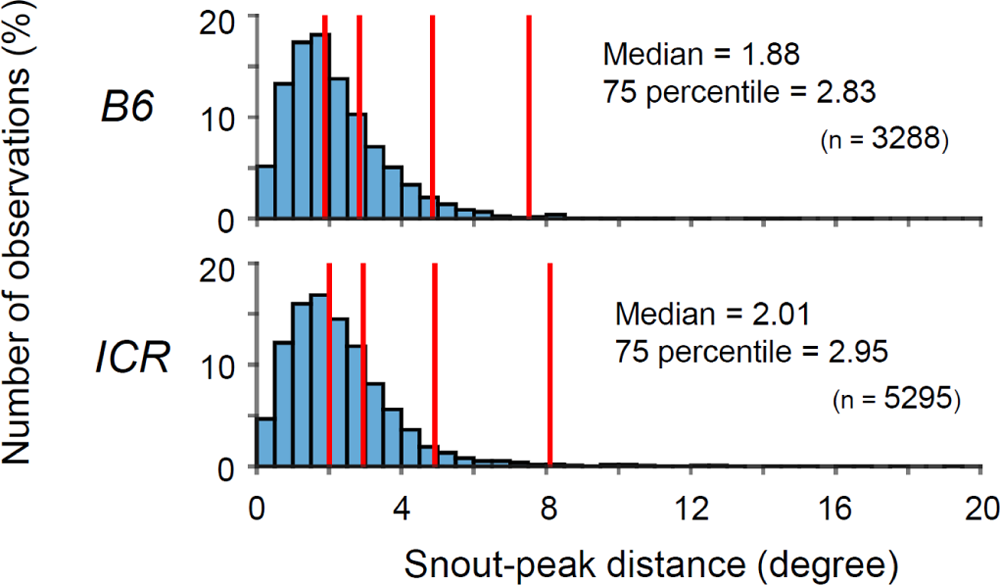
Error histograms similar to those in Figure 1e, calculated separately for B6 (top) and ICR (bottom) mice.

**Supplementary Figure 3.**
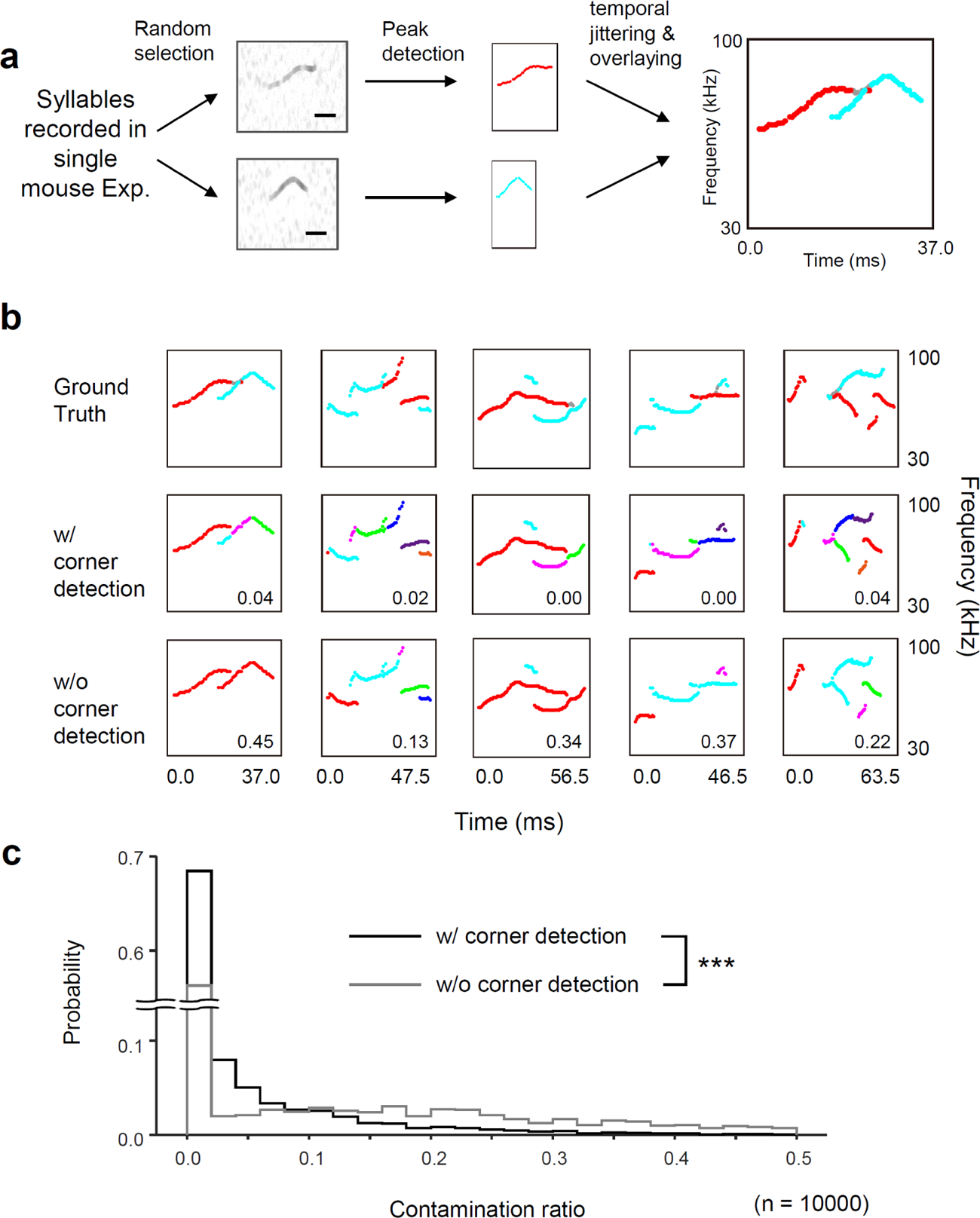
Effectiveness of the modified segmentation algorithm. (**a**) Test data generation. Scale bars in the spectrograms = 20 ms. Gray peaks were excluded to simulate the peak search in the USVSEG algorithm. (**b**) Examples of the ground truth labels (top) and segmentation with (middle) and without (bottom) corner detection (Supplementary Figure 1e) for separation at the crossing points. Different colors of the ground truth indicate different sources. Different colors in the segmentation results indicate different segments. The number in each segmentation result indicates the contamination ratio. (**c**) The distribution of contamination ratios in the segmentation with (black) and without (gray) corner detection. \*\*\**p* = 1.9 × 10^-117^, two-tailed Wilcoxon rank-sum test.

**Supplementary Figure 4.**
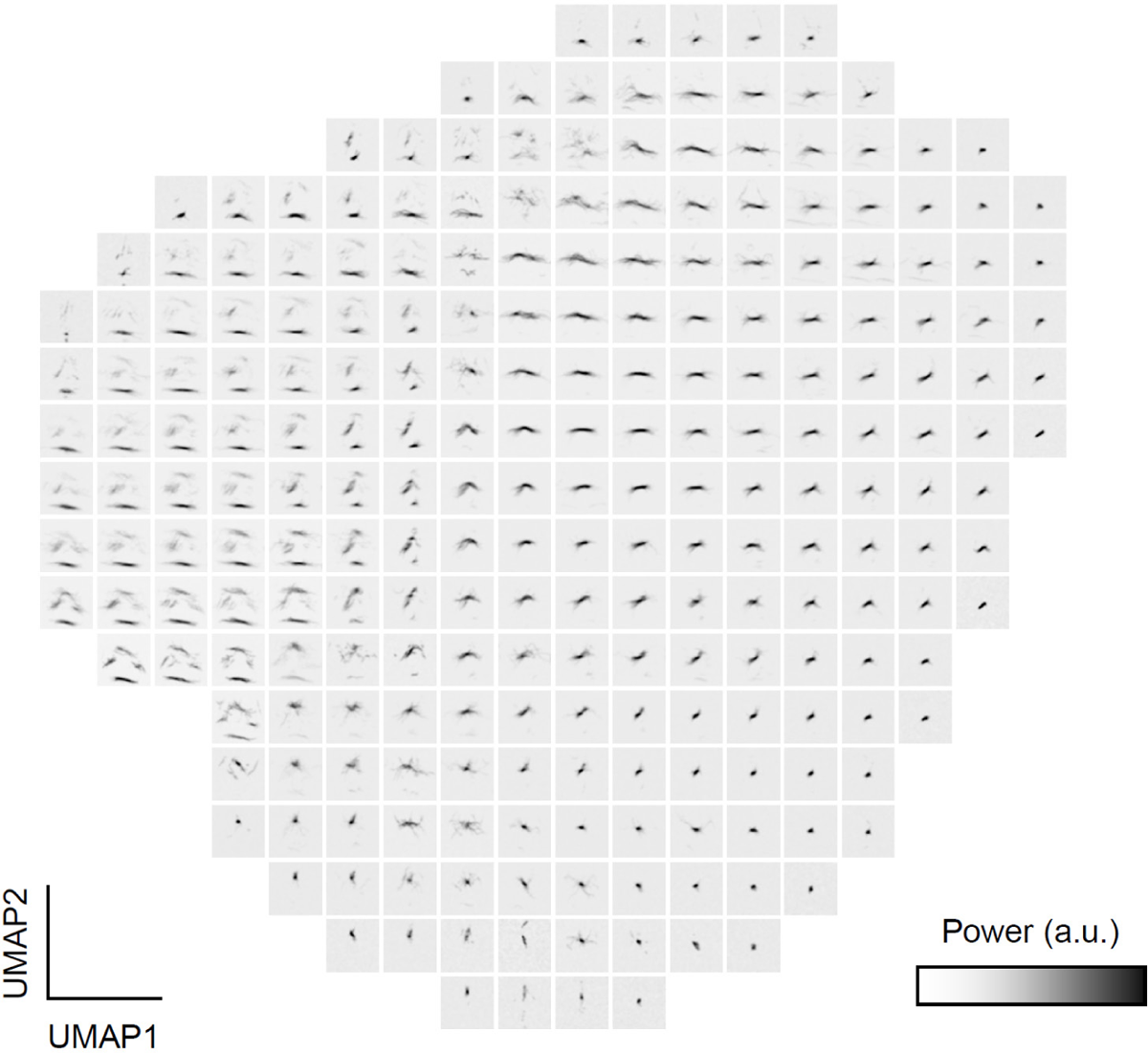
Mean spectrograms of the syllables for each location on the UMAP projection in Fig. 2c. The frequency range of the spectrograms is 30 to 100 kHz, and the time range is 128 ms.

**Supplementary Figure 5.**
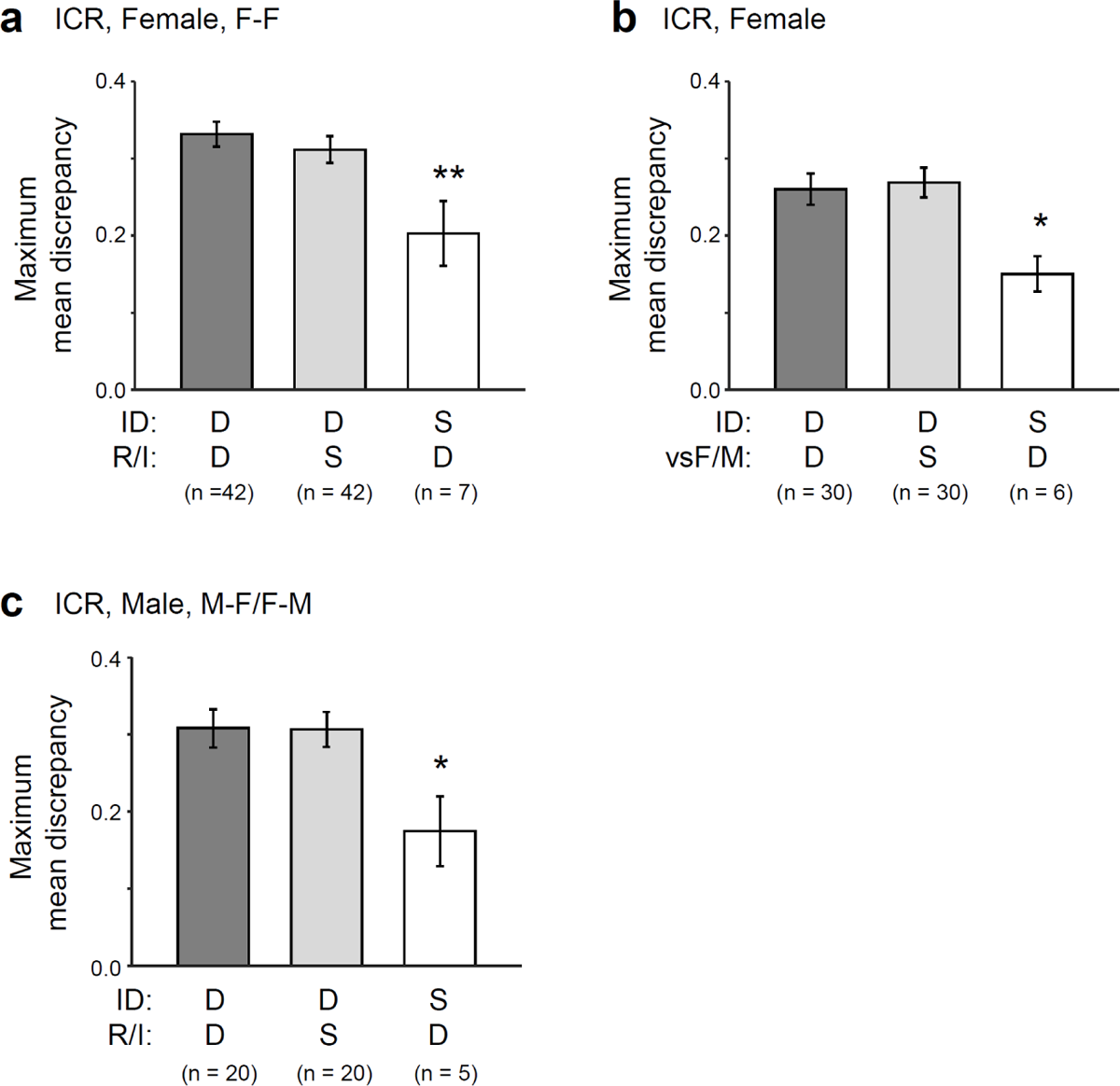
Quantitative analysis of the similarity of vocal repertoires within the same social contexts or individuals. Dissimilarities in vocal repertoire calculated as maximum mean discrepancy (MMD) between pairs of distributions of the features of syllables (Fig. 2c) emitted by individuals. D, different; S, same. Error bars, s.e.m. (**a**) Comparisons of mean MMDs between groups of pairs of different IDs and different R/I labels (dark gray), pairs of different IDs and the same R/I labels (light gray), and pairs of the same IDs and different R/I labels (white) in female ICR mice in F-F sessions. \*\**p* < 0.01, difference from pairs of different IDs and different R/I labels (dark gray) with Dunnett’s test after one-way analysis of variance (ANOVA) (F[2,88] = 4.154; *p* = 0.0189). (**b**) Comparison of mean MMDs between groups of pairs of different IDs and different vsF/M labels (dark gray), pairs of different IDs and the same vsF/M labels (light gray), and pairs of the same IDs and different vsF/M labels (white) in female ICR mice in F-F, F-M, and M-F sessions. \**p* < 0.05, difference from pairs of different IDs and different vsF/M labels (dark gray) with Dunnett’s test after one-way ANOVA (F[2,63] = 3.261; *p* = 0.0449). (**c**) Similar comparison as (**a**) in male ICR mice in F-M and M-F sessions. \**p*<0.05, difference from pairs of different IDs and different R/I labels (dark gray) with Dunnett’s test after one-way ANOVA (F[2,42] = 3.504; *p* = 0.0391).

**Supplementary Figure 6.**
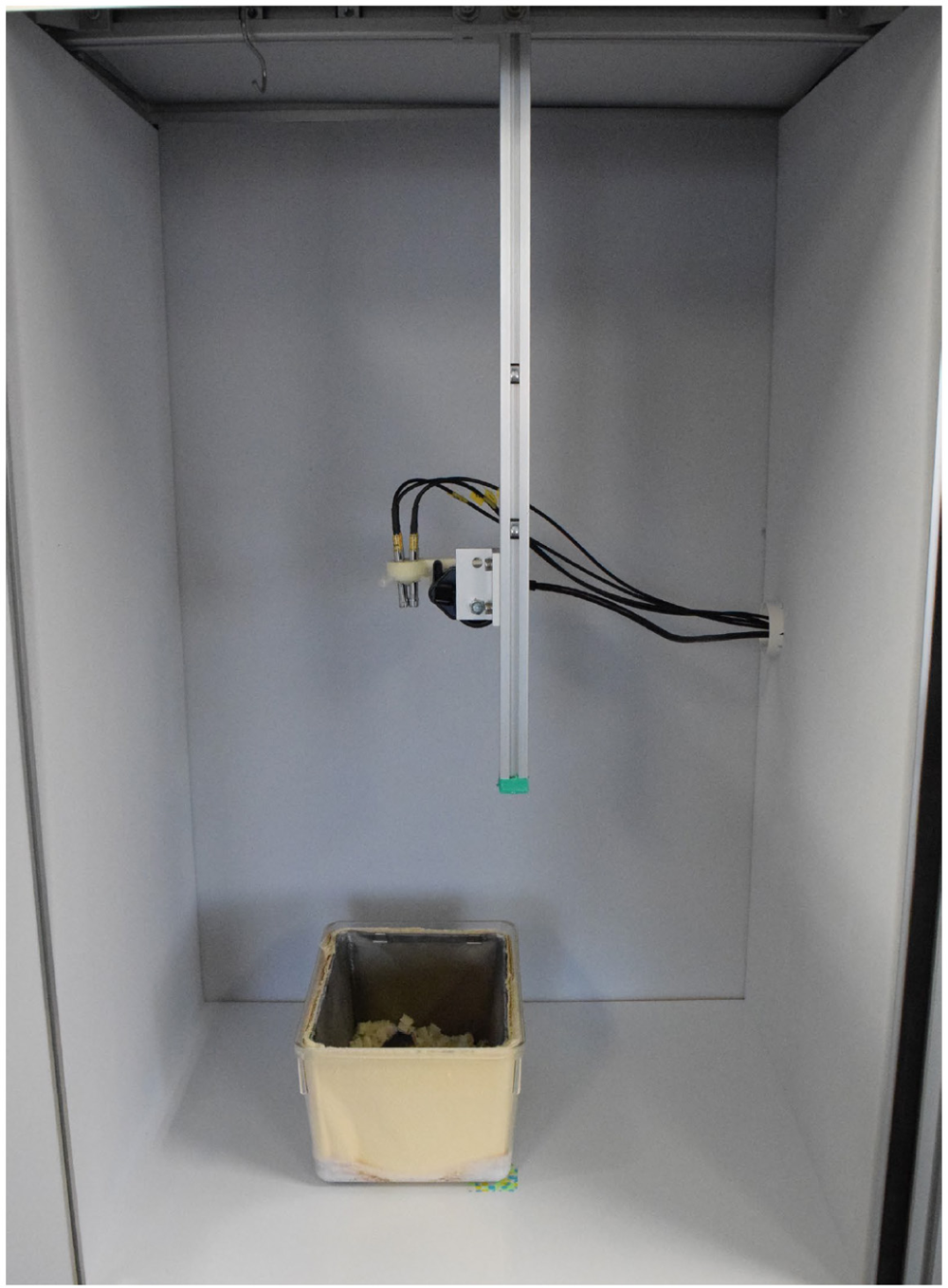
The soundproof box used in this study. The floor was made of polyvinyl chloride. The walls and ceiling were covered with sound-absorbing melamine foam to reduce sound reflections.

**Supplementary Figure 7.**
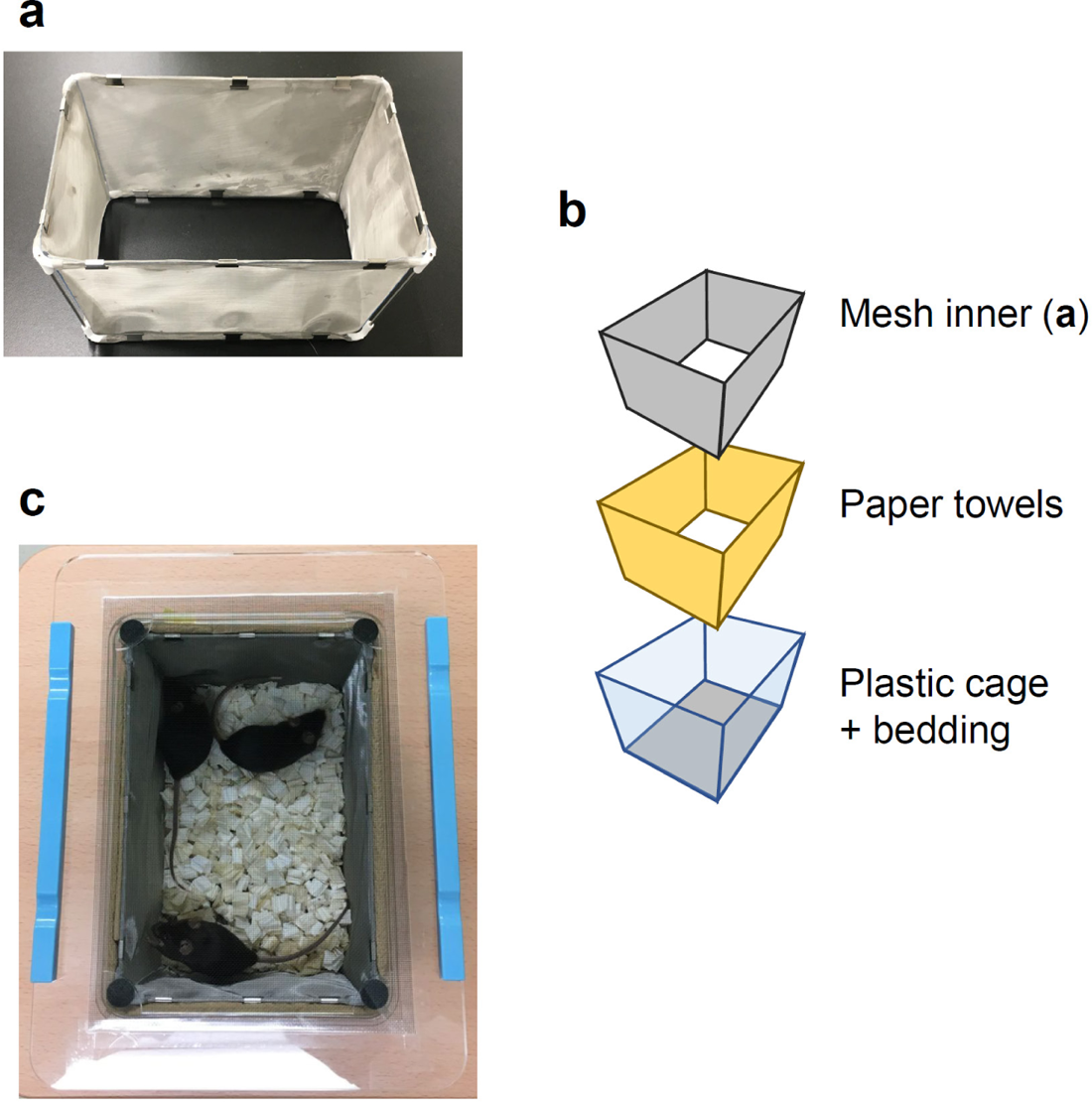
Details of the home cage recording. (**a**) The inner cage was made of fine stainless mesh. (**b**) An illustration of the layers of the cage. (**c**) The cage lid used for recording. A clear mesh screen was glued onto an acrylic frame. Rubber feet were attached to the corners of the lid to pin down the inner cage. The light blue bars are weights.

**Supplementary Figure 8.**
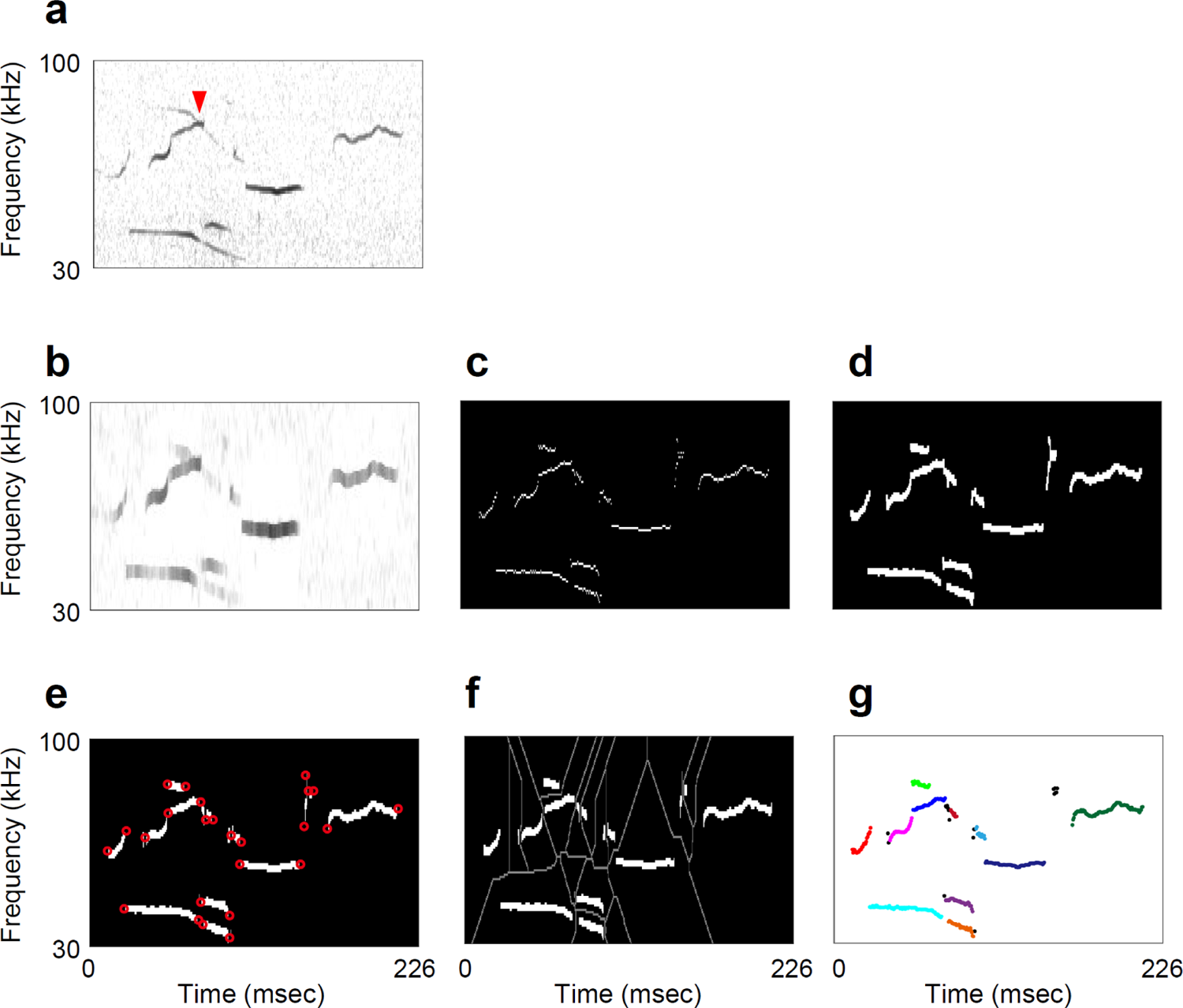
USV segmentation. **(a)** A spectrogram of example USVs. Note that in this example, calls emitted from two different mice crossed at the red arrow. **(b–g)** The segmentation process. First, a filtered spectrogram was calculated **(b)**, and peaks in the spectrogram were extracted **(c)** using the USVSEG algorithm. Then, morphological dilation and erosion were repeatedly applied to connect the continuous peaks **(d)**. To separate the segments at the crossing points, corners of the images were detected and removed (**e**; red circles indicate the detected corners). Finally, the watershed algorithm was applied to find the boundary of the segments (**f**; gray curves indicate the boundaries of the segments). The resultant segments are shown in **g**. The colors represent different segments. Segments that were too short (≤ 3 ms; colored black) were excluded from subsequent localization processes.

**Supplementary Figure 9.**
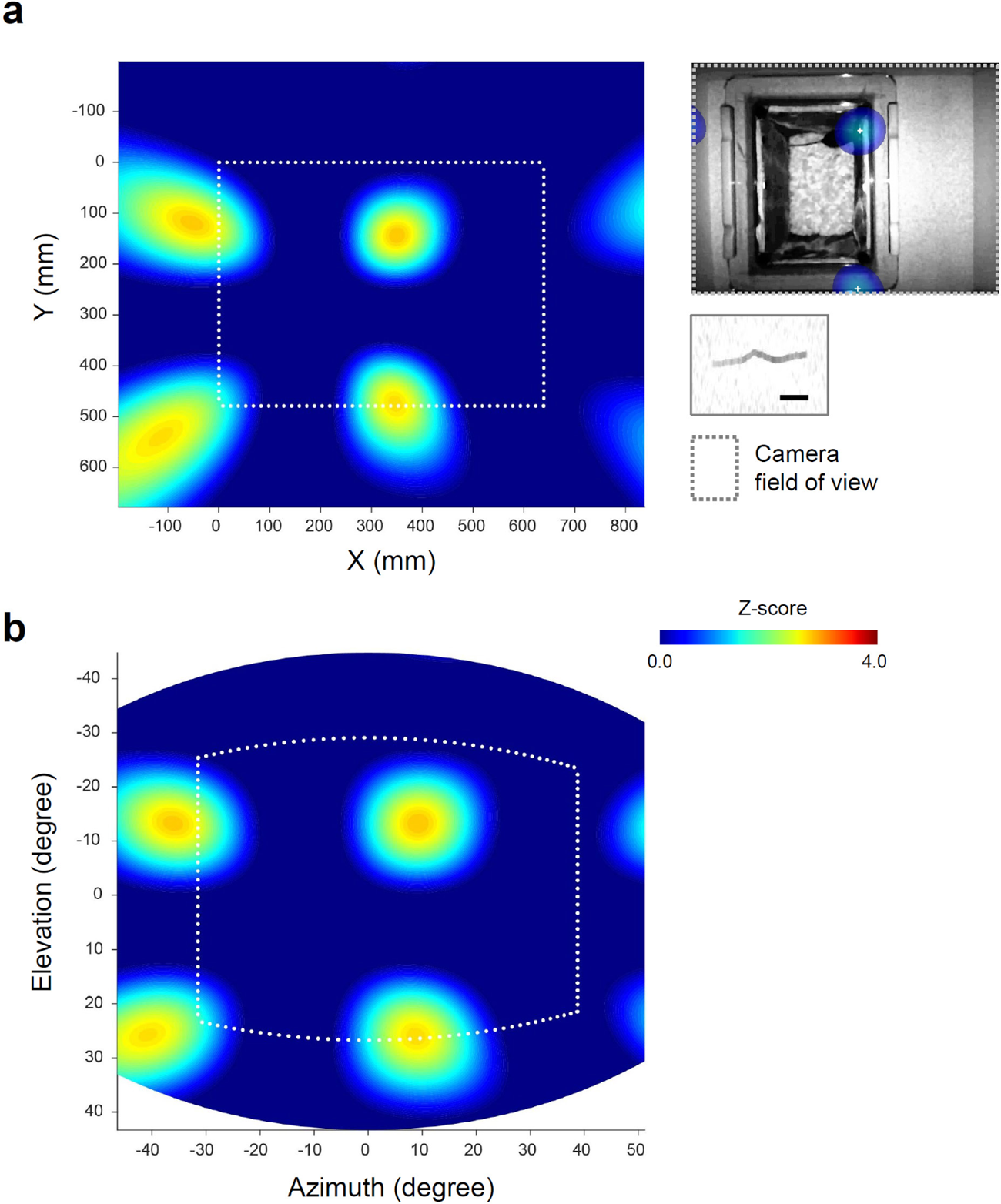
An example of the spatial spectrum shown in a wide field of view. (**a**) Spatial spectrum around the camera field of view. Inset shows the spatial spectrum overlayed on the video frame (top) and sound spectrogram (frequency range = 30 to 100 kHz, scale bar = 20 ms) of a USV segment. (**b**) The same spatial spectrum projected in the direction from the center of the microphone array. The peaks are located in a square grid.

**Supplementary Figure 10.**
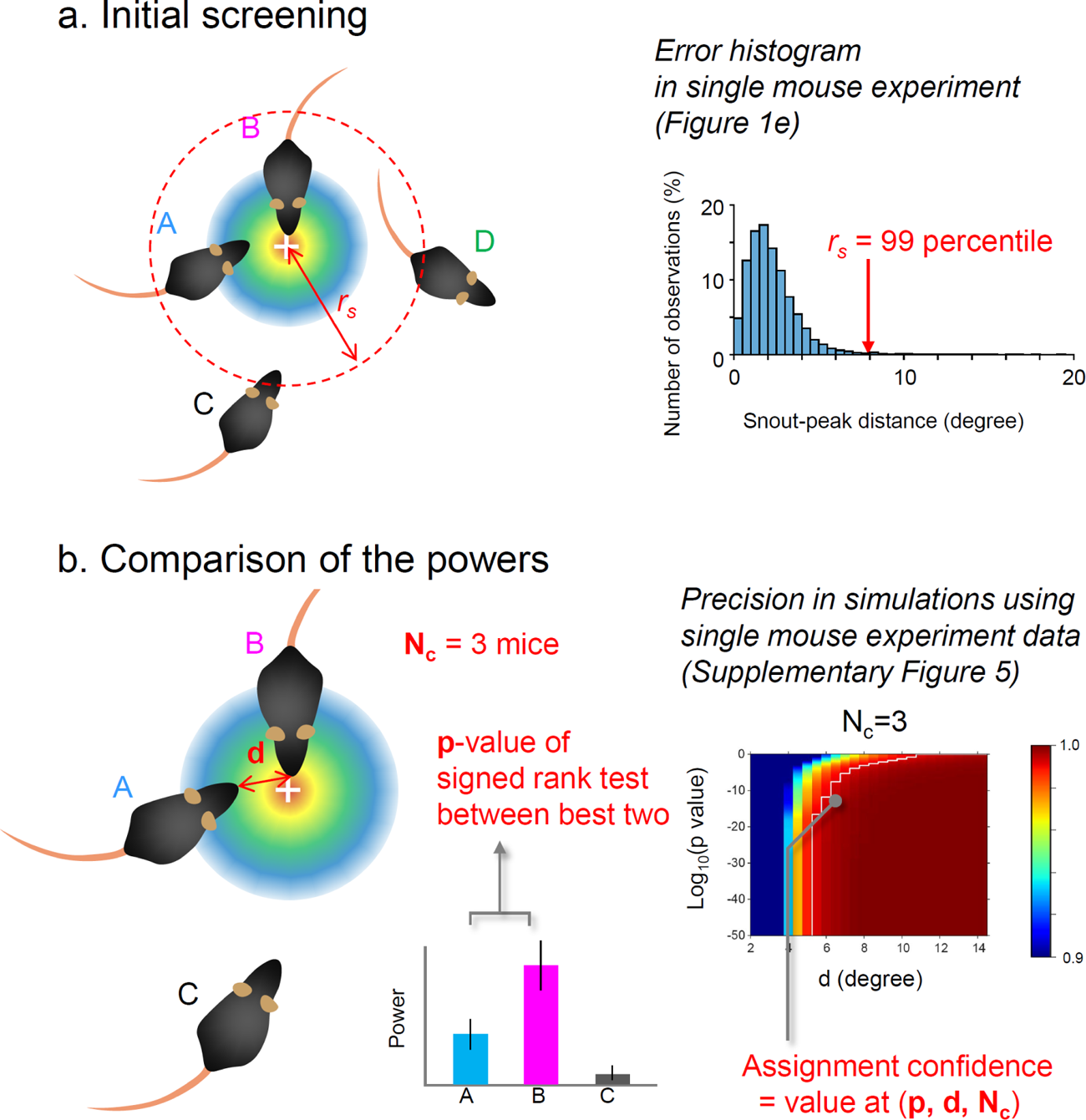
USV assignment. Assignment was performed in two steps: initial screening. (**a**) and comparison of the sound powers at the snouts of mice (**b**). (**a**) First, mice with peak-snout distances that were longer than the threshold (r_s_) were excluded from the candidates. A–D, IDs of mice. The white cross indicates the peak of the spatial spectrogram. (**b**) The USV segment was then assigned to the mouse with the highest power if the assignment confidence was higher than 99%. The confidence value was estimated using the *p*-value of the comparison between the best two mice (*p*), the distance between the snouts of the best two mice (*d*), and the number of candidate mice (N_c_), based on simulations using the data of the single mouse experiments. See Methods for details.

**Supplementary Figure 11.**
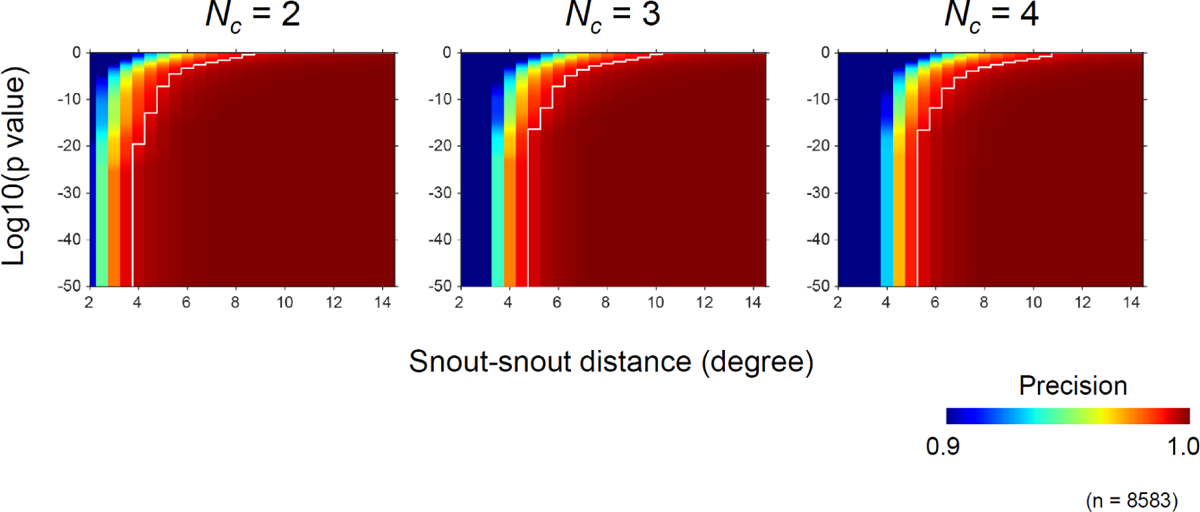
Distribution of assignment precision in the simulation using data obtained from the single mouse experiments. p, *p*-value of the sign-rank test comparing the sound powers at the snouts between the best two mice. *N_c_*, the number of candidate mice. White curves indicate the border of 0.99.

**Supplementary Figure 12.**
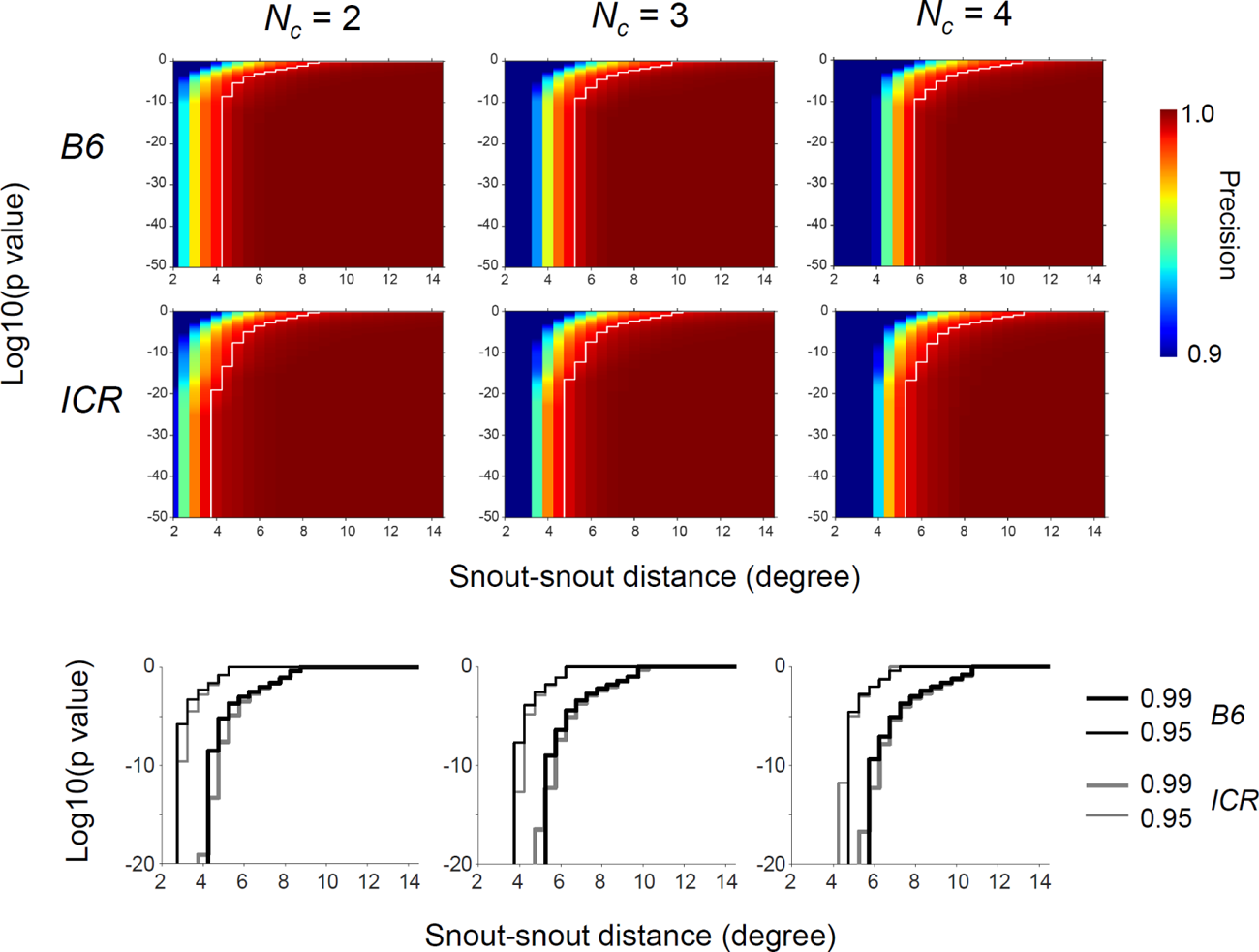
Distribution of assignment precision similar to that in Supplementary Figure 11, calculated separately for B6 and ICR mice. Top: distribution of the precision. Bottom, comparison of the borders of 0.95 and 0.99 between strains. Because the simulation results for B6 and ICR mice were largely similar to those above, we used a combined distribution (Supplementary Fig. 11) for the assignments in this study.

**Supplementary Figure 13.**
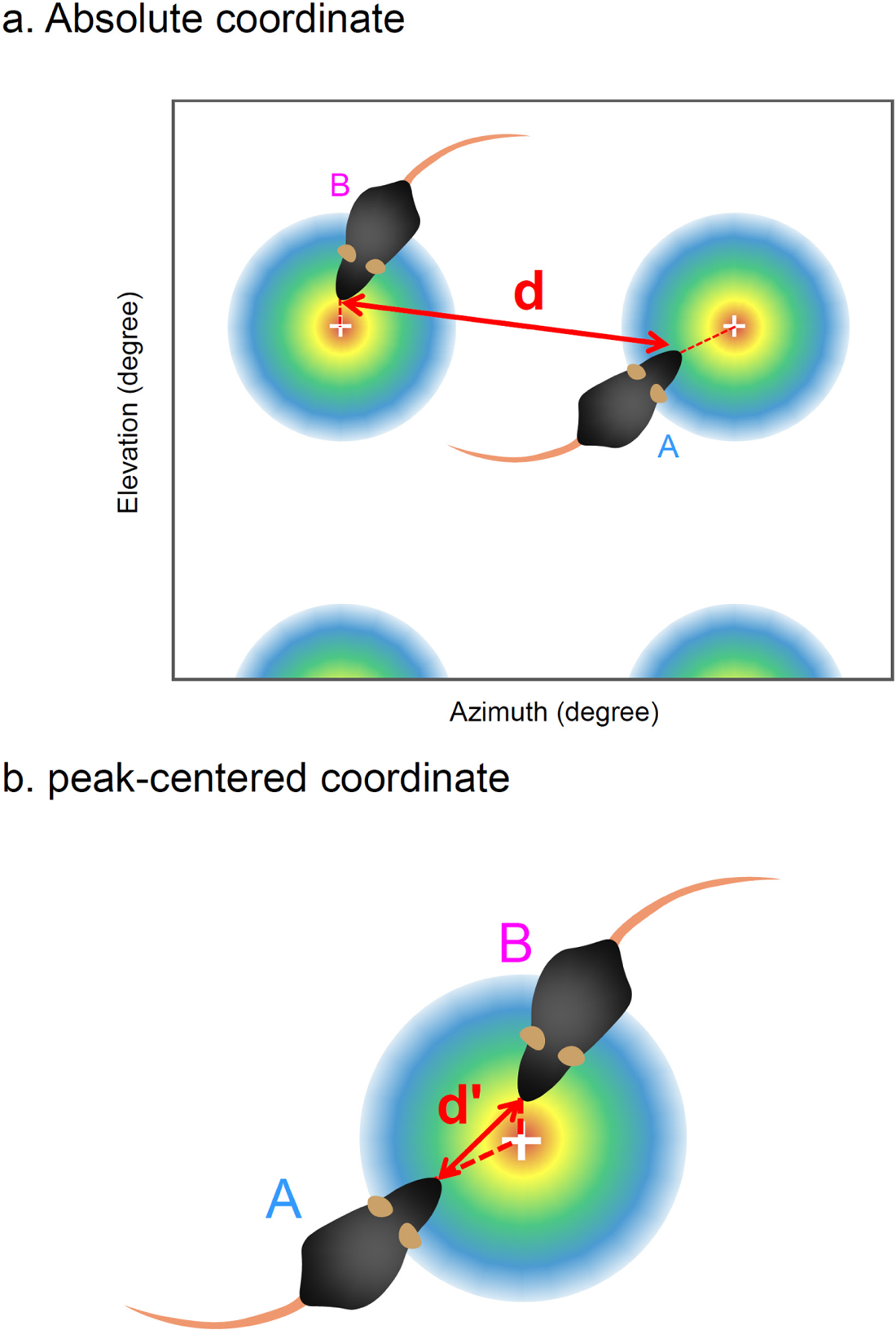
Spatial ‘wrapping’ for fixing snout-snout distance for the assignment. **a**) An illustration of a case where mice are near different peaks. (**b**) The mouse locations in **a** are aligned to the peak-centered coordinates. The snout-snout distance in this space (*d’*) was used for estimating assignment confidence. A and B, IDs of mice.

**Supplementary Figure 14.**
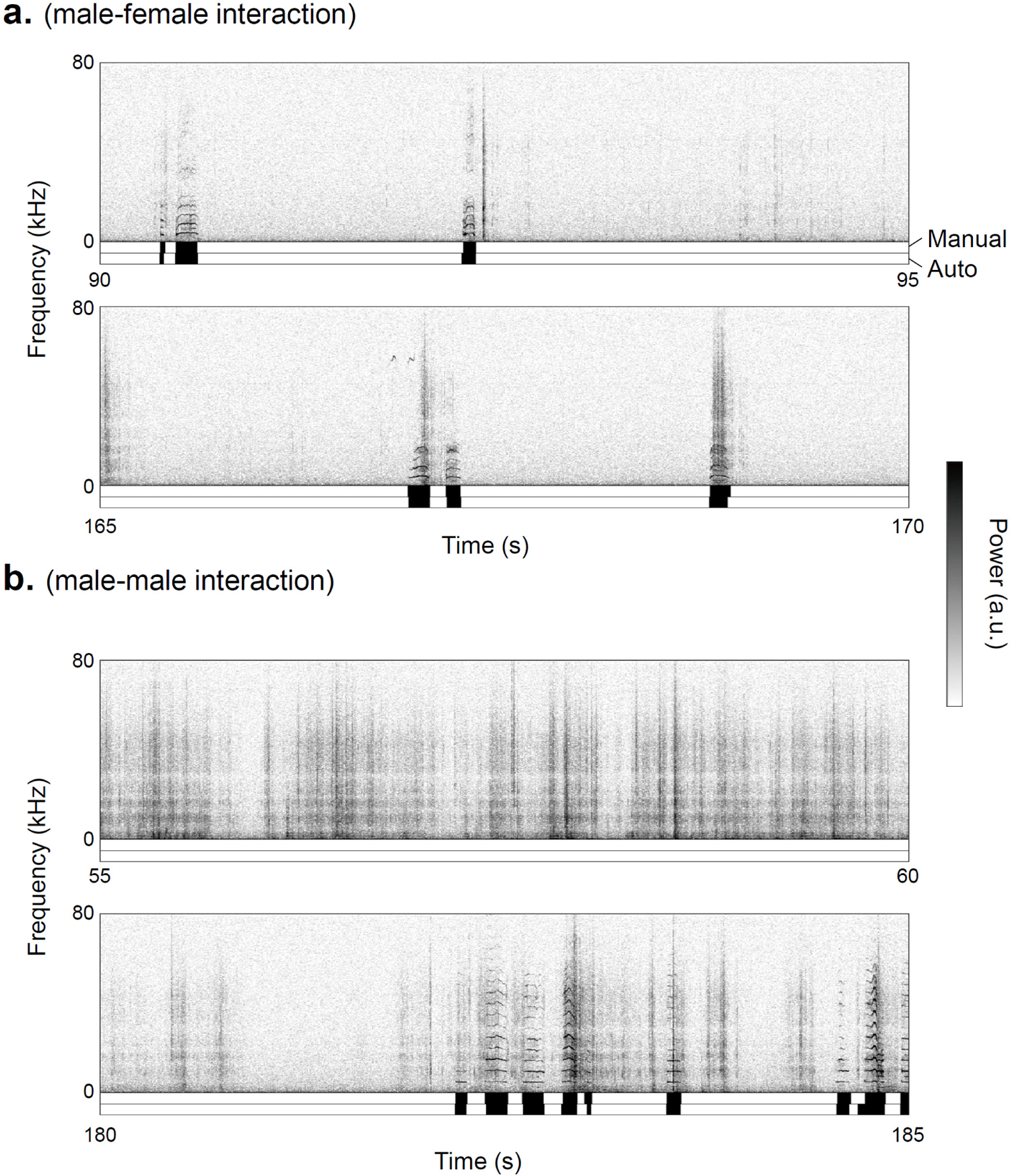
Examples of BBV detection using the automatic detector. Examples of 5 s spectrograms are shown in male-female (**a**) and male-male (**b**) ICR mice interactions. Black bars in the first and second rows below the spectrogram indicate manual annotation and automatic detection of BBVs, respectively.

**Supplementary Figure 15.**
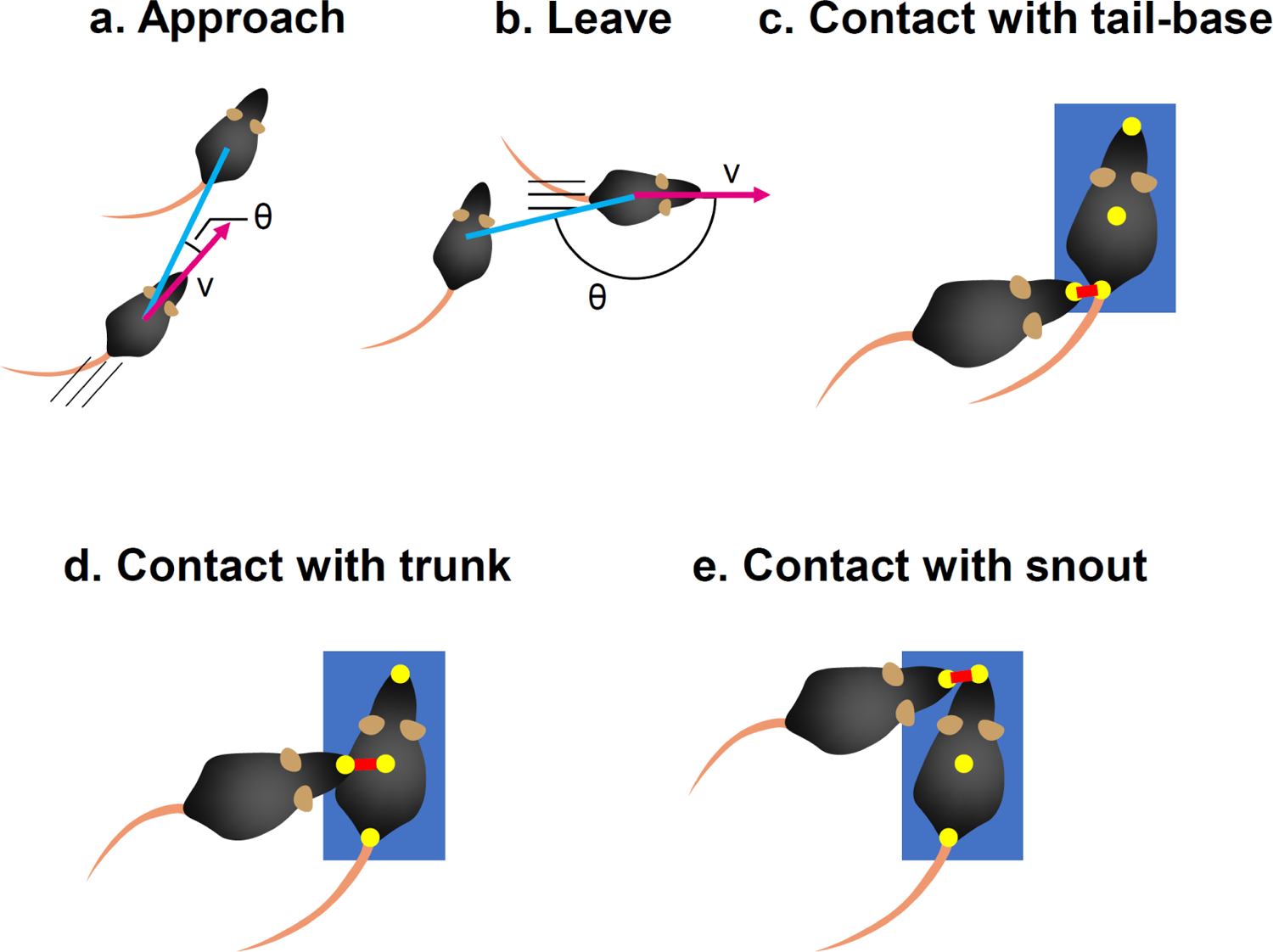
Definitions of the behavioral events. *Approach* (**a**) was counted when a subject’s running speed (v) was > 50 mm/s, and the angle between the partner location and running direction (θ) was < 45 degrees. *Leave* (**b**) was counted when v was > 50 mm and θ was > 135 degrees. *Contact with the tail-base* (**c**), *trunk* (**d**), and *snout* (**e**) was counted when a subject’s snout was inside the bounding box of the partner (blue shaded area), and the body part (yellow point) of the partner closest to subject’s snout was the tail-base, the center of the bounding box, and the snout, respectively. The trunk position was calculated as the center of the bounding box.

**Supplementary Table 1.** Assignment performance in the simulations based on the single mouse experiment.

|  |  |  | Assigned |  |  | Unassigned |  |  |
| --- | --- | --- | --- | --- | --- | --- | --- | --- |
|  | Sub. | Total | Hit | Error | Precision | Low Conf. | Unlocalized | Pos. N/A |
| B6 | 2 | 3384 | 2873<br>(84.9%) | 5<br>(0.1%) | **99.8%** | 410<br>(12.1%) | 65<br>(1.9%) | 31<br>(0.9%) |
|  | 3 | 3384 | 2537<br>(75.0%) | 15<br>(0.4%) | **99.4%** | 736<br>(21.7%) | 65<br>(1.9%) | 31<br>(0.9%) |
|  | 4 | 3384 | 2213<br>(65.4%) | 23<br>(0.7%) | **99.0%** | 1052<br>(31.1%) | 65<br>(1.9%) | 31<br>(0.9%) |
| ICR | 2 | 5678 | 4777<br>(84.1%) | 17<br>(0.3%) | **99.6%** | 501<br>(8.8%) | 293<br>(5.2%) | 90<br>(1.6%) |
|  | 3 | 5678 | 4347<br>(76.6%) | 32<br>(0.6%) | **99.3%** | 916<br>(16.1%) | 293<br>(5.2%) | 90<br>(1.6%) |
|  | 4 | 5678 | 3956<br>(69.7%) | 31<br>(0.5%) | **99.2%** | 1308<br>(23.0%) | 293<br>(5.2%) | 90<br>(1.6%) |

Sub., number of subjects in the cage; one is the real mouse and the others are virtual mice randomly positioned in the cage.
Total, total number of USV segments detected.
Hit, number of segments correctly assigned to the real mouse.
Error, number of segments incorrectly assigned to one of the virtual mice.
Precision, percentage of Hit ÷ (Hit + Error).
Low Conf., number of segments in which assignment confidences were < 0.99.
Unlocalized, number of unlocalized segments.
Pos. N/A, number of segments emitted when the snout positions could not be estimated because of video-based tracking issues.
Audible call, number of segments that overlapped with audible calls.

**Supplementary Table 2.** Assignment performance in the home cage social interaction experiment.

|  |  |  |  | Unassigned |  |  |
| --- | --- | --- | --- | --- | --- | --- |
|  | Sub. | Total | Assigned | Low Conf. | Unlocalized | Pos. N/A |
| B6 | 2<br>(F-F) | 102 | 84<br>(82.4%) | 14<br>(13.7%) | 2<br>(2.0%) | 2<br>(2.0%) |
|  | 2<br>(M-M) | 30 | 26<br>(86.7%) | 3<br>(10.0%) | 0<br>(0.0%) | 1<br>(3.3%) |
|  | 2<br>(F-M) | 1795 | 1420<br>(79.1%) | 339<br>(18.9%) | 14<br>(0.8%) | 22<br>(1.2%) |
|  | 2<br>(M-F) | 2518 | 1823<br>(72.4%) | 483<br>(19.2%) | 29<br>(1.2%) | 183<br>(7.3%) |
|  | 3<br>(M-FF) | 970 | 578<br>(59.6%) | 263<br>(27.1%) | 52<br>(5.4%) | 77<br>(7.9%) |
| ICR | 2<br>(F-F) | 16052 | 9813<br>(61.1%) | 3203<br>(20.0%) | 468<br>(2.9%) | 2568<br>(16.0%) |
|  | 2<br>(M-M) | 743 | 191<br>(25.7%) | 288<br>(38.8%) | 27<br>(3.6%) | 237<br>(31.9%) |
|  | 2<br>(F-M) | 8030 | 3881<br>(48.3%) | 1760<br>(21.9%) | 236<br>(2.9%) | 2153<br>(26.8%) |
|  | 2<br>(M-F) | 2897 | 1497<br>(51.7%) | 759<br>(26.2%) | 109<br>(3.8%) | 532<br>(18.4%) |
|  | 3<br>(M-FF) | 1607 | 569<br>(35.4%) | 804<br>(50.0%) | 69<br>(4.3%) | 165<br>(10.3%) |

Sub., number of subjects in the cage; the first and second letters in the parentheses indicate resident and intruder(s). M, one male; F, one female; FF, two females. Total, total number of USV segments detected. Assigned, number of segments assigned to one of the mice. Low Conf., number of segments in which assignment confidences were < 0.99. Unlocalized, number of unlocalized segments. Pos. N/A, number of segments emitted when the snout positions could not be estimated because of video-based tracking issues. Percentages in parentheses represent ratios relative to the total number of segments.

**Supplementary Table 3.**
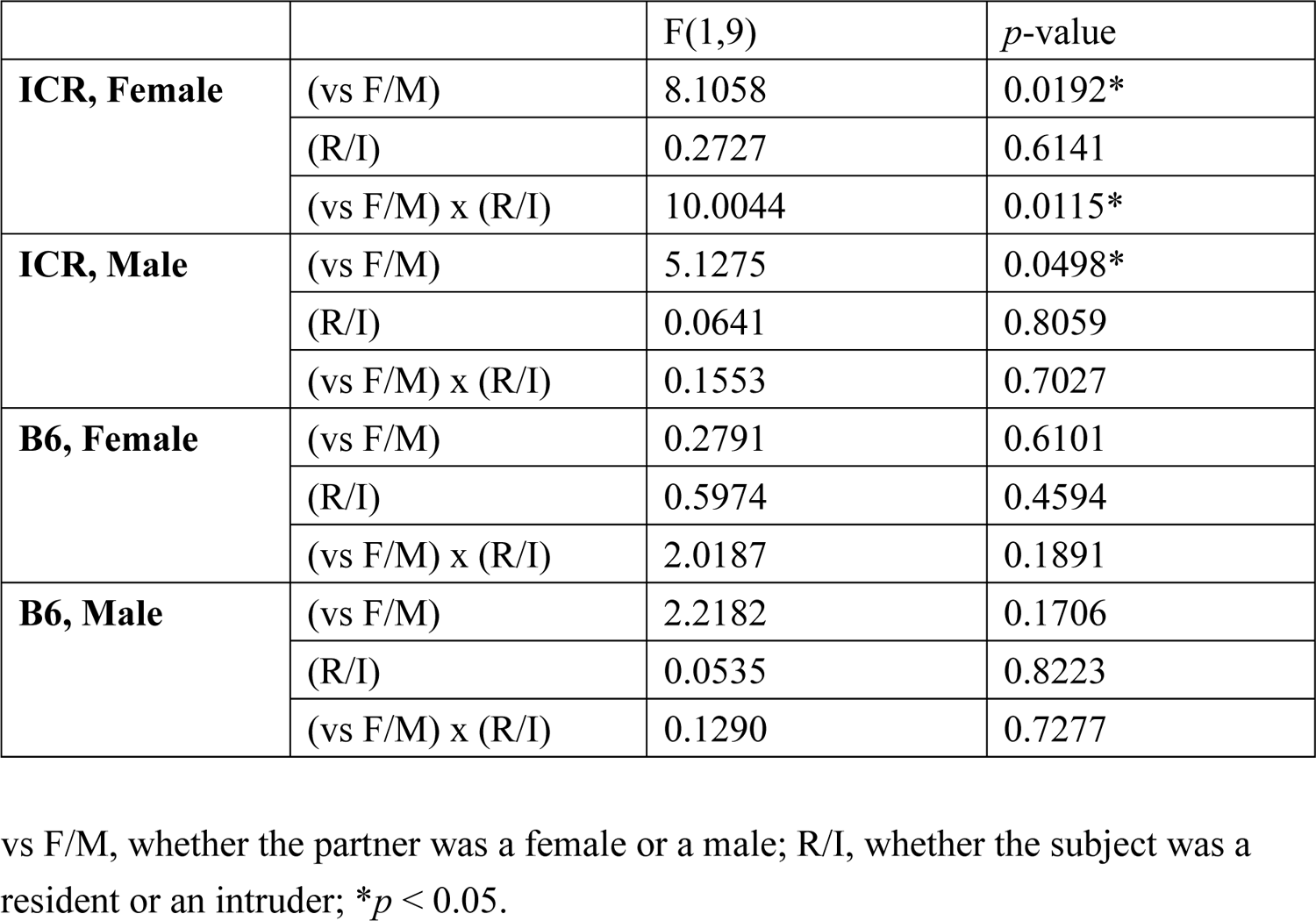
Detailed results of the two-way ANOVA in Fig. 2a.

**Supplementary Table 4.**
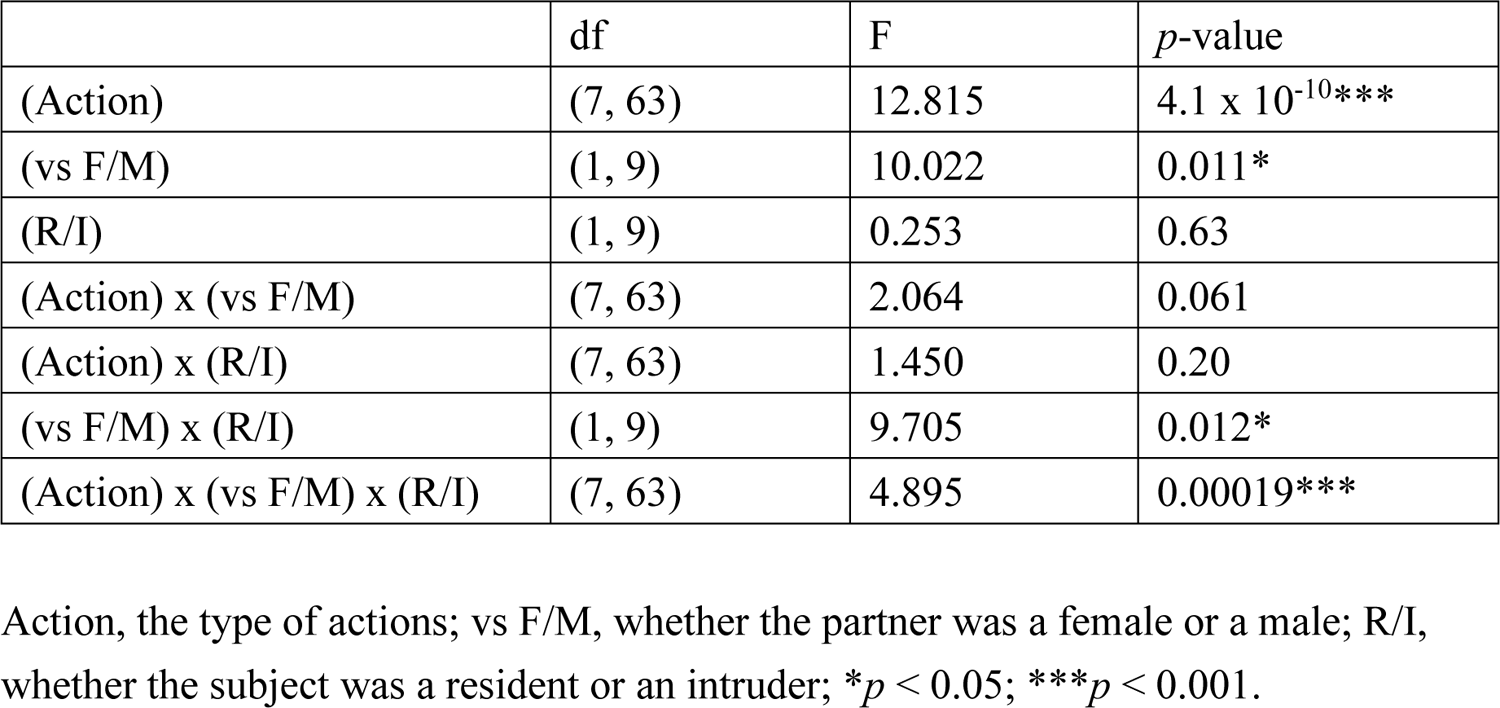
Detailed results of the three-way ANOVA in Fig. 2b.

**Supplementary Table 5.** Number of ultrasound segments that overlapped with BBVs.

|  | Sub. | # of segments |
| --- | --- | --- |
| B6 | 2<br>(F-F) | 0 |
|  | 2<br>(M-M) | 1 |
|  | 2<br>(F-M) | 0 |
|  | 2<br>(M-F) | 3 |
|  | 3<br>(M-FF) | 0 |
| ICR | 2<br>(F-F) | 99 |
|  | 2<br>(M-M) | 959 |
|  | 2<br>(F-M) | 834 |
|  | 2<br>(M-F) | 1999 |
|  | 3<br>(M-FF) | 1236 |

Sub., number of subjects in the cage; the first and second letters in parentheses indicate resident and intruder(s). M, one male; F, one female; FF, two females.

**Supplementary Video 1.** An example of USV localizations emitted from a male B6 mouse in a home cage of a female mouse.

**Supplementary Video 2.** An example of USV localizations emitted from a male ICR mouse in a home cage of a female mouse.

**Supplementary Video 3.** An example of USV assignment during an interaction between a male (cyan) and a female (magenta) B6 mouse in a home cage.

**Supplementary Video 4.** An example of USV assignment during an interaction between two female ICR mice in a home cage.

**Supplementary Video 5.** An example of the video tracking result used for behavioral event classification.

